# Estimating Boltzmann Statistical Energy for Proteins from 3D Structure Using Pairwise Amino Acid Energy Matrix - GEM-X2120-N871

**DOI:** 10.1101/2025.11.17.688978

**Authors:** Kunchur Guruprasad

## Abstract

A pairwise amino acid energy matrix: GEM-X2120-N871 is developed for evaluating the Boltzmann statistical energy (BSE) for proteins of known 3D structure. The matrix was derived from a dataset of representative protein crystal structures and NMR structures selected from the Protein Data Bank (PDB). The pairwise amino acid contacts separated by distance ≤ 3.2 Å in protein 3D structures was used to evaluate probabilities and propensities against background and the Boltzmann equation *P(x) = e^-(E(x)/KT)^* that links the probability of observing an event *x* with energy *E(x)* at a given temperature *T* was used to evaluate the BSE for proteins defined in Millielectron Volts (MeV) units. The BSE for a protein is obtained by summing up corresponding values for amino acid pairs from the matrix that satisfy the distance criterion in protein 3D structure. The landscape of BSE values for representative proteins based on propensities ranged between −890.99 MeV to 711.57 MeV among crystal structures and between −717.64 MeV to 234.78 MeV among NMR structures. Whereas, the landscape of BSE values based on probability alone was comparatively much narrower and ranged between 36.88 MeV to 131.28 MeV for the crystal structures and between 34.30 MeV to 102.23 MeV for the 871 NMR structures. Therefore, BSEs based on propensity with a broader range of values is suggested as being useful to discriminate individual proteins. The evaluation of BSE for proteins can be used to estimate relative gain or loss in energy between proteins that has applications in identifying low-energy conformations among an ensemble of protein conformations, or changes in energy due to drug/inhibitor/small molecule binding to the protein. The method was applied to evaluate changes in BSE for certain cancer drugs bound to protein targets and to analyse trends in predictions based on BSE with experimental observations on drug/inhibitor binding effectiveness as reported in literature.

## 1. Introduction

Statistical models based upon evaluation of propensities and free-energy functions derived from frequency of occurrence of structural arrangement of residues/atoms in a collection of protein three-dimensional structures can be used to estimate the relative strengths of the protein chain interactions (Shortle, 2003). The frequencies of occurrences of structural contacts can be converted into an estimate of its free energy based upon the Boltzmann hypothesis, for instance, hydrogen bonds, hydrophobicity, side-chain interactions, proline isomerization, amino acid phi/psi propensities have been shown to conform to the Boltzmann hypothesis (Shortle, 2003 and references therein). The Boltzmann’s equation *P(x) = e^-(E(x)/KT)^* defines the probability of observing an event *x* with energy *E(x)* at a given temperature *T*. Hence, the energy of an event observed with a probability *P(x)* can be computed as *E(x)* = -*KT* ln[*P(x)*], thereby, higher the energy of an event (with respect to *KT*) is, lower is its probability.

The Protein Data Bank (PDB) (Berman et al., 2000) (https://www.rcsb.org), a central data repository for experimentally determined protein three-dimensional structures, contains the Cartesian co-ordinates (x, y, z) corresponding to the atomic positions of amino acid residues labelled ‘ATOM’ in the PDB file. The PDB file also contains co-ordinates for other atomic positions, labelled ‘HETATM’ that correspond either to ligands, substrates, peptides, inhibitors, metal ions, co-factors, prosthetic groups, glycosylation, DNA, RNA, drugs, inhibitors, sugars, solvent molecules or other such moieties defined in protein three-dimensional structure. Thereby, the atomic positions corresponding to amino acid residues resulting from the accommodation of different moieties in protein three-dimensional structures can be used to evaluate the residue-pair probability/propensity and corresponding Boltzmann statistical energy according to the Boltzmann’s equation.

In this work, the Boltzmann statistical energies (BSE) were evaluated using the Boltzmann’s equation based upon the probability and propensity values for pairwise amino acid contacts separated by a distance ≤ 3.2 Å against background in protein three-dimensional structures. Thereby, amino acids distant along the sequence but close in protein three-dimensional structure are taken into account as they would contribute to the pairwise interaction energy associated with the amino acids. The GEM-X2120-N871 matrix proposed contains weights derived for 400 amino acid pairs based upon the analysis of a total of 2991 representative protein three-dimensional structures in the Protein Data Bank (PDB). The landscape of the BSE values for proteins in Millielectron Volts (MeV) was examined. The method was applied for evaluating the BSEs for certain cancer drug-target protein complexes. This study facilitated assessment of the observed trends in BSE prediction for proteins with drug inhibition studies based upon experimental reports in literature.

## 2. Materials and Methods

The representative protein chains corresponding to the protein three-dimensional structures in PDB were identified according to the PDB_SELECT program (Hobohm et al., 1992). Accordingly, non-redundant protein chains that shared < 25% sequence identity among protein crystal structures determined at ≤ 2.5 Å resolution and the NMR (Model_1) structures were selected from the PDB (Berman et al., 2000) (https://www.rcsb.org). Protein chains comprising amino acid residues labelled ‘X’ and the obsolete entries were excluded. The atom coordinates corresponding to amino acid residues labelled ‘ATOM’ were extracted from the individual PDB files. The amino acid pairs within contact distance ≤ 3.2 Å between non-hydrogen atoms were identified and the probability P(x) values for residue-pair contacts in above range normalized against background were evaluated. The BSE was evaluated using the Boltzmann’s equation E(x)=-kTln[P(x)] defined in Millielectron Volts per Kelvin units and the value of kT at temperature 298K as 25.7 MeV was used to evaluate the BSE. Further, the propensity values were evaluated for the amino acid pairs against background by taking ratio of the probability values, i.e., number of a particular amino acid pair ≤ 3.2 Å in all proteins divided by the total number of all amino acid pairs ≤ 3.2 Å in the dataset and this value divided by the ratio of the number of particular amino acid pair in all proteins divided by the total number of all amino acid pairs in the dataset. The BSE for amino acid pairs represented the weights in GEM-X2120-N871 matrix. The matrix was used for evaluating the BSE for protein by summing up the individual weight values corresponding to amino acid pairs separated by distance ≤ 3.2 Å in protein three-dimensional structure. The computer programs for analyses were developed in FORTRAN programming language at ABREAST™ (https://www.abreast.in). All figures were generated using Microsoft Excel software.

## 3. Results and discussions

The protein structures selected for analyses comprised: 1) 2120 protein chain crystal structures and 2) 871 NMR structures. The PDB codes corresponding to the protein crystal structures are attached in **Supplementary Data-A** and for the NMR structures in **Supplementary Data-B.** The proteins sequences ranged between 19 to 636 amino acid residues with ∼62.11% protein chains comprising < 100 amino acid residues, ∼36.94% comprising between 101-400 amino acid residues and ∼0.93% comprising between 400 to 636 amino acid residues. The analysis dataset contained a total of 2991 protein chains that comprised 558,524 amino acid pairs with ≤ 3.2 Å contacts against a background of 27,622,539 amino acid pairs.

### 3.1. Pairwise Amino Acid Propensities and the Boltzmann statistical pairwise amino acid energy (weight) matrix

The propensity values for 400 amino acid pairs are shown in **Figure 1**. The ‘CC’ pair showed a distinctively high propensity value of 5.72 compared to the other amino acid pairs and the ‘PE’ pair was associated with the least propensity 0.47. The Boltzmann statistical energy values (or weights) for the amino acid pairs are shown in **Figure 2**. According to the Boltzmann’s equation, the energy values are inversely correlated to the propensity values, i.e., higher the propensity, lower is the Boltzmann statistical pairwise amino acid energy and vice-versa. Accordingly, the ‘CC’ amino acid pair is associated with least energy −44.85 MeV and the ‘PE’ amino acid pair is associated with maximum energy value 19.12 MeV. The BSE values (weights) applicable to experimentally determined protein three-dimensional structures based upon crystal structures and NMR structures and referred as ‘GEM-X2120-N871’ matrix is shown in **Table 1**. The amino acids listed along rows represents the first amino acid in the pair and the amino acids listed along the columns represents the second amino acid in the pair. It was observed that 214 out of 400 amino acid pairs had negative energy values, suggesting an almost equal distribution of the amino acid pairs associated with negative and positive energy values and therefore a good representation of the protein dataset is used for the analyses. The amino acid pairs associated with low energy values (< −9.0 MeV) were: CC, HH, HC, KE, RE, KD, EK, RC, CY, GC, KC, CW, RD, ER and the amino acid pairs with relatively higher energy values, i.e., (> 9.0 MeV) mainly comprised proline (‘P’) as first residue in the pair as shown in **Figure 2**.

**Figure 1.**
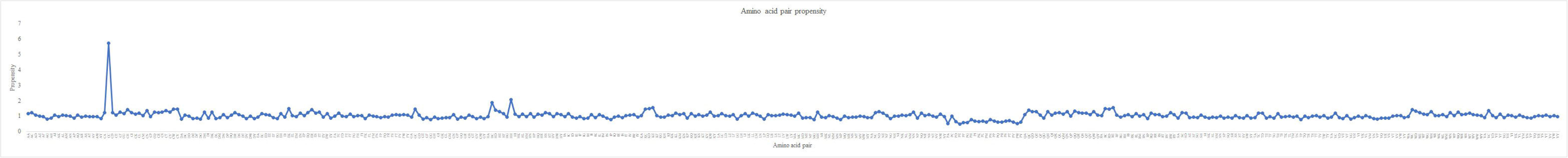
Pairwise amino acid propensity values for the 400 amino acid pairs.

**Figure 2.**
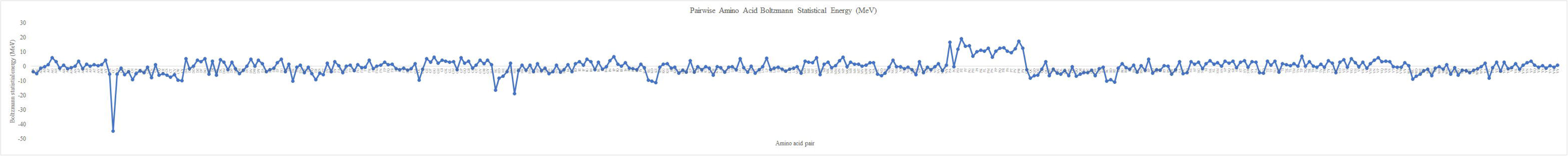
Boltzmann statistical pairwise amino acid energy values (in MeV) for the 400 amino acid pairs.

**Table 1.**
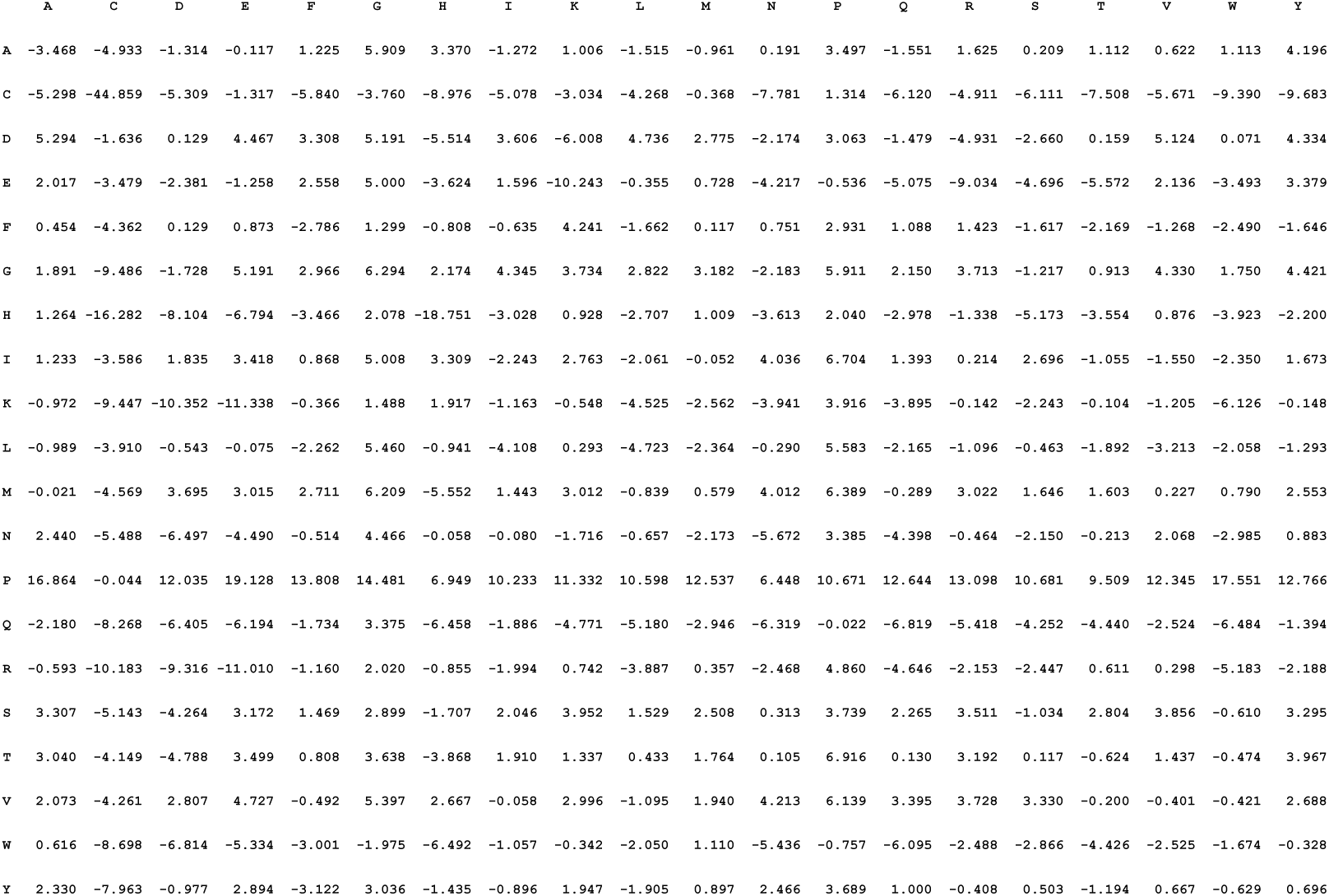
Boltzmann statistical pairwise amino acid energy matrix: GEM-X2120-N871.

### 3.2. Boltzmann statistical energy landscapes for representative protein crystal structures and NMR structures in the Protein Data Bank

The Boltzmann statistical energy values evaluated using the GEM-X2120-N871 matrix (as in **Table 1**) for the 2120 protein crystal structures are attached in **Supplementary Data-C** and for the 871 protein NMR structures in **Supplementary Data-D**. These Supplementary data contain information on protein PDB code/chain, total number of amino acid residues in protein chain, total number of amino acid residue pairwise contacts (≤ 3.2 Å) in the protein chain three-dimensional structure and the computed BSE values (in MeV units). The landscape of BSE values ranged between −890.99 MeV to 711.57 MeV for crystal structures (refer **Supplementary Data-C**) and between −717.64 MeV to 234.78 MeV for the 871 protein NMR structures (refer **Supplementary Data-D**). However, majority of the proteins in the combined dataset were associated with BSEs that lie in range between −250.0 MeV to +250 MeV. The BSEs calculated based on probabilities for the proteins ranged between 36.88 MeV to 131.28 MeV for the 2120 protein crystal structures and between 34.30 MeV to 102.23 MeV for the 871 NMR structures. Thereby, evaluation of BSEs for protein three-dimensional structures based on propensities rather than probabilities gives a broader range of BSE values that facilitates ranking of the proteins based on energy. Accordingly, the proteins associated with low BSEs were: integrin beta-2 (PDB code:2P28B), copper storage protein (csp3) (5ARMA), 25 kda ookinete surface antigen (6B0GE), phospholipase A2 (1G4IA) and the proteins associated with relatively high BSEs were: FprA *Mycobacterium tuberculosis* oxidoreductase (1LQTB), fructose-1,6-bisphosphate aldolase/phosphatase (3T2CA), cytochrome c551 peroxidase (4AANA), Alnumycin P phosphatase AlnB (4EX6A), glutamate-1-semialdehyde 2,1-aminomutase from *Arabidopsis thaliana* (5HDMA). The success of the Boltzmann hypothesis in explaining the energetics of protein structures (as applicable to real proteins in the biological context) has been attributed to evolutionary dynamics where protein sequences and structures interact over time to maintain stability (Shortle, 2003).

### 3.3. Assessing the Boltzmann statistical energy values for some of the cancer drug-target protein complexes in the Protein Data Bank

The Boltzmann statistical energy was evaluated for some of the cancer drug target protein complexes available in the PDB. The crystal structures of the Bcr-Abl kinase domain and certain mutants bound to drugs/inhibitors were analysed. The drugs represented small molecule inhibitors of the Bcr-Abl kinase domain used in the treatment of chronic myelogenous leukemia (CML). The crystal structures represented kinase domain of c-Abl in complex with the small molecule inhibitors; imatinib drug (STI-571/Gleevec) (PDB code:1IEP) and PD173955 (Parke-Davis) (PDB code:1M52) (Nagar et al., 2002), crystal structure of dasatinib drug (BMS-354825) bound to activated Abl kinase domain (PDB code:2GQG) (Tokarski et al., 2006), crystal structure of T315I mutant in complex with Aurora kinase PHA-739358 inhibitor (PDB code:2V7A) (Modugno et al., 2007), crystal structure of human Abl kinase in complex with nilotinib (AMN-107) (PDB code:3CS9) (Weisberg et al., 2005), crystal structure of Abl kinase domain bound with DFG-out inhibitor ponatinib (AP24534) that inhibits both native and mutant Bcr-Abl including T315I acting as pan-Bcr-Abl inhibitor (PDB code: 3OXZ)) (O’Hare et al., 2009, Zhou et al., 2011) and the crystal structure of the ABL T315I mutant kinase domain bound with a DFG-out inhibitor AP24589 (PDB code: 3OY3) (Zhou et al., 2011). The BSEs for these proteins are shown in **Figure 5**.

**Figure 3.**
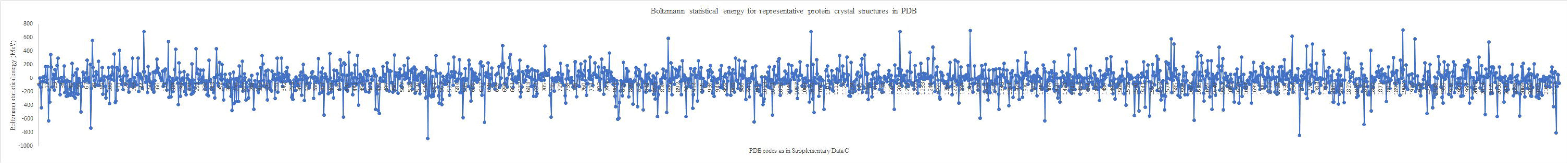
Landscape of the Boltzmann statistical energy (BSE) values (in MeV) units based on GEM-X2120-N871 pairwise amino acid energy matrix for 2120 representative crystal structures in the Protein Data Bank.

**Figure 4.**
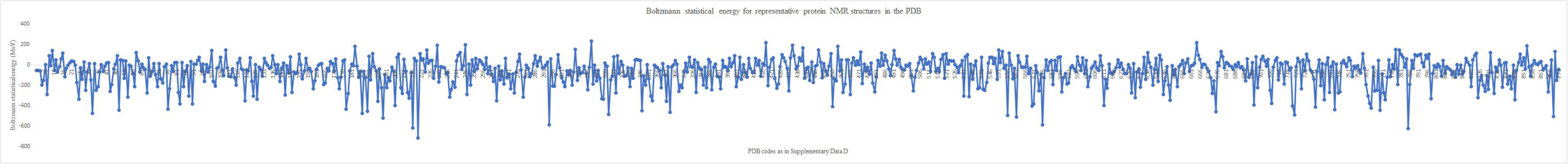
Landscape of the Boltzmann statistical energy (BSE) values (in MeV) units based on GEM-X2120-N871 pairwise amino acid energy matrix for 871 representative NMR structures in the Protein Data Bank.

**Figure 5.**
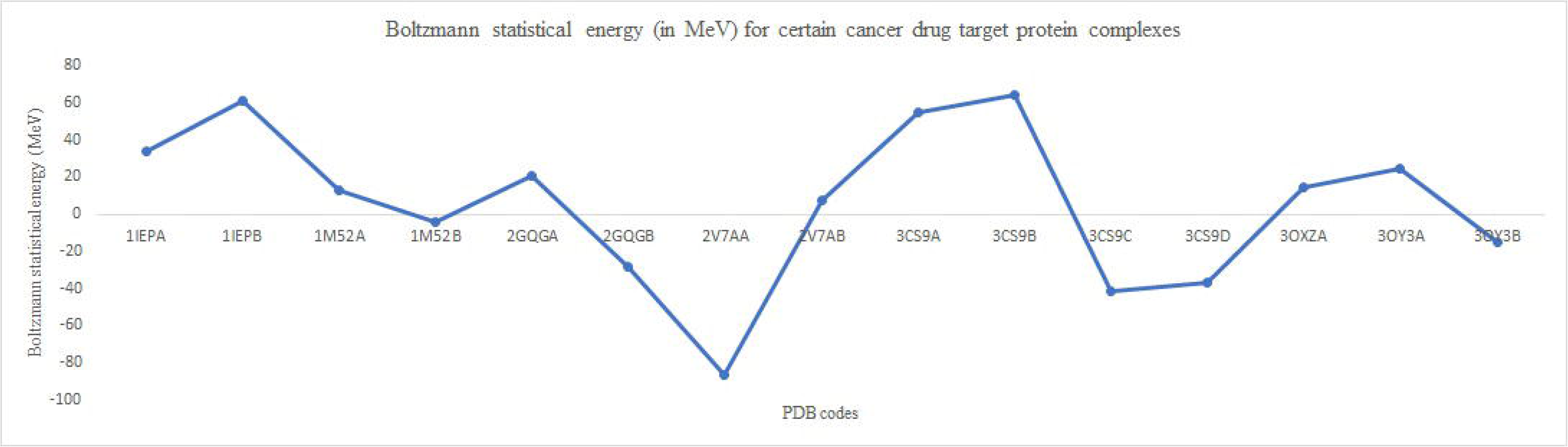
Boltzmann statistical energy values evaluated using the GEM-X2120-N871 pairwise amino acid energy matrix for some of the cancer drug target protein complexes in the PDB.

The imatinib bound Bcr-Abl kinase domain complex was associated with BSE 33.77 MeV (PDB code:1IEP_A) and 61.31 MeV (PDB code:1IEP_B). The chains comprised 274 amino acid residues each but with different number of contacts; A-chain (563) and B-chain (559). The imatinib bound to the protein A_chain was associated with low BSE. The binding of PD173955 to Bcr-Abl was associated with relatively low BSE values; 12.75 MeV (PDB code:1M52_A) and −4.06 MeV (PDB code:1M52_B). Thereby, comparing BSE values associated with PD173955 and that of imatinib binding to Bcr-Abl, relatively low energy values were associated with PD173955 binding. This observation is consistent with the *in-vitro* kinase assays that showed 10-fold more inhibitory activity of PD173955 compared to that of imatinib (Nagar et al., 2002). In the case of PD173955 binding to the protein target, the B-chain was associated with comparatively low energy.

Dasatinib (BMS-354825), a multi-targeted tyrosine kinase inhibitor targeting the oncogenic pathways, is known to be a more potent inhibitor than imatinib against wild-type Bcr-Abl (Tokarski et al., 2006). The BSE values were; 20.63 MeV (PDB code: 2GQG_A) and −28.71 MeV (PDB code: 2GQG_B) that is relatively low compared to equivalent values observed for imatinib binding (PDB code:1IEP). This observation is consistent with the finding by (Tokarski et al., 2006) that dasatinib is more potent than imatinib in binding to the Bcr-Abl tyrosine kinase domain. The B-chain was associated with low energy. The resistance to therapy with imatinib in patients with chronic myelogenous leukemia (CML) mainly due to the mutations in Bcr-Abl led to the development of second generation inhibitors that were able to overcome most imatinib-resistant mutants, except the T315I mutation in Bcr-Abl.

The Aurora kinase inhibitor, PHA-739358 inhibited *in-vitro* the kinase activity of Bcr-Abl and several mutants including the T315I mutation (Modugno et al., 2007). The co-crystal structure complex of the Aurora kinase inhibitor, PHA-739358 with the T315I mutant Abl kinase (PDB code:2V7A) was used to compute the BSE. Accordingly, the BSE values were; −86.71 MeV (PDB code:2V7A_A) and 7.76 MeV (PDB code:2V7A_B). These values were relatively low compared to the equivalent values observed for imatinib binding to the Bcr-Abl. In retrospect, the results of the prediction using the GEM-X2120-N871 matrix support choice of PHA-739358 as a candidate molecule for validation targeting the Bcr-Abl.

Another known selective inhibitor of Bcr-Abl, recognized as a promising inhibitor for the Bcr-Abl tyrosine kinase oncogene that causes CML and Philadelphia chromosome-positive (Ph+) acute lymphoblastic leukemia (ALL) was that of AMN109 (nilotinib). This inhibitor is significantly more potent than imatinib and active against number of imatinib-resistant Bcr-Abl mutants and its three-dimensional crystal structure is known (PDB code:3CS9) (Weisberg et al., 2005). The BSE values computed for the inhibitor bound to the different protein chains in the structure were; 55 MeV (PDB code: 3CS9_A), 64.1 MeV (B-chain), −41.19 MeV (C-chain), −36.75 MeV (D-chain). The C-chain was associated with relatively low energy value compared to imatinib (STI-571 or Gleevec) or dasatinib (BMS-354825) bound to the Bcr-Abl kinase target, thereby supporting further development of nilotinib (AMN107) inhibitors in drug discovery program for therapy of CML, ALL.

Another compound, AP24534 (ponatinib) bound to the Bcr-Abl kinase domain as in the crystal structure of (PDB code: 3OXZ) (Zhou et al., 2011) provided insights into how the ponatinib binds to the ABL kinase domain, including its interactions with key residues that help overcome resistant mutations, such as, T315I mutation that is useful to develop new inhibitors against emerging compound mutations. The BSE values for AP24534 (PDB code: 3OXZ) were; 14.71 MeV (A-chain) and for AP24589 were 24.8 MeV (PDB code: 3OY3, A-chain) and −14.89 MeV (PDB code: 3OY3, B-chain). These values are relatively low compared to the values for imatinib binding supporting development of these compounds in the pan Bcr-Abl inhibitor drug discovery programs for treatment of CML. The pairwise amino acid contact energy matrix GEM-X2120-N871 and the method developed to evaluate BSE for proteins of known three-dimensional structure can be used to facilitate choice of potential compounds for design and validation in drug discovery programs.

### 3.4. Discussions

In summary, a matrix: GEM-X2120-N871 corresponding to the BSE values (or weights) for the 400 amino acid pairs is proposed. The matrix is derived from contact propensities for amino acid pairs separated by distance ≤ 3.2 Å against background among representative protein three-dimensional structures selected from the PDB. The distance threshold ensures selection of amino acid pairs capable of forming hydrogen-bonds involving their main-chain or side-chain atoms. Accordingly, the BSEs for protein is estimated in Millielectron Volts (MeV) by summing up the corresponding values derived from the matrix for amino acid pairs satisfying the distance criterion in protein three-dimensional structure. The GEM matrix and method developed can be applied to estimate the relative gain or loss in energy between proteins, to identify low energy conformers among ensemble of protein conformations and to select appropriate protein chain templates for lead optimization in structure-based drug discovery programs.

## 4. Conclusions

A computational method has been developed for evaluating the Boltzmann Statistical Energy (BSE) in Millielectron Volts (MeV) units for proteins of known three-dimensional structure. The method is based on evaluation of BSEs using the Boltzmann’s equation for propensities of pairwise amino acid contacts separated by distance ≤ 3.2 Å against background among representative crystal structures and NMR structures selected from the PDB. A matrix of 400 pairwise amino acid statistical energy values called the GEM-X2120-N871 matrix is proposed. The landscape of BSE values evaluated for the representative proteins in the PDB lie between −890.99 MeV to 711.57 MeV. The method applied to evaluate the BSEs for some of the cancer drug-target protein complexes showed agreement in trends between the predictions based on energy from this work and drug/inhibitor binding effectiveness based on experiments as reported in the literature. The GEM-X2120-N871 BSE matrix of pairwise amino acid energy values can be used to evaluate the BSE for proteins of known three-dimensional structure that can have applications in studies related to estimating the relative changes in energy between proteins and in drug discovery programs.

## Supplementary Data

**Supplementary Data-A.** The Protein Data Bank (PDB) codes corresponding to 2120 representative protein crystal structures.

**Supplementary Data-B.** The Protein Data Bank (PDB) codes corresponding to 871 representative protein NMR structures.

**Supplementary Data-C.** The Boltzmann statistical energy values (in MeV) evaluated for the 2120 protein crystal structures using GEM-X2120-N871 matrix.

**Supplementary Data-D.** The Boltzmann statistical energy values (in MeV) evaluated for the 871 protein NMR structures using GEM-X2120-N871 matrix.

## CRediT authorship contribution statement

**Kunchur Guruprasad:** Conceived the project, developed computer programs in FORTRAN programming language required for the work, carried out analyses and wrote the manuscript.

## Acknowledgements

The author gratefully acknowledges the Protein Data Bank used in the research work. The computer programs for the analyses were developed at ABREAST™ (https://www.abreast.in).

## Data availability statement

All data is available in the manuscript and as Supplementary data.

