## Supplementary-Data-C for "Estimating Boltzmann Statistical Energy for Proteins from 3D Structure Using Pairwise Amino Acid Energy Matrix - GEM-X2120-N871"

| PDB | No of aa's | No of ints. | X2120-N871 dataset |
|---|---|---|---|
| 1A2XB | 31 | 79 | -94.64 |
| 1A73A | 162 | 282 | 2.49 |
| 1AH7A | 245 | 530 | -134.92 |
| 1ALUA | 157 | 325 | -438.68 |
| 1ASSA | 152 | 309 | -49.69 |
| 1AVFQ | 22 | 37 | 21.76 |
| 1B0NB | 31 | 58 | 11.07 |
| 1B33O | 67 | 115 | -49.81 |
| 1B35D | 57 | 92 | 23.69 |
| 1BAVC | 307 | 631 | 174.77 |
| 1BKRA | 108 | 209 | -61.72 |
| 1BM8A | 99 | 204 | -22.84 |
| 1BQCA | 302 | 648 | 171.12 |
| 1BTEB | 94 | 148 | -629.79 |
| 1BV4D | 109 | 194 | -173.28 |
| 1BX7A | 51 | 77 | -353.66 |
| 1C0PA | 363 | 674 | 350.25 |
| 1C5EC | 95 | 158 | 166.46 |
| 1CBNA | 46 | 86 | -124.76 |
| 1CDLG | 20 | 43 | -15.52 |
| 1CKSC | 78 | 106 | 59.49 |
| 1CLVI | 32 | 54 | -173.18 |
| 1CN3F | 29 | 43 | 123.74 |
| 1CQ4A | 37 | 59 | 34.21 |
| 1CQ4B | 23 | 33 | -1.78 |
| 1CSEI | 63 | 114 | 189.06 |
| 1CYWA | 159 | 302 | 296.55 |
| 1CZBA | 141 | 288 | -9.62 |
| 1CZTA | 160 | 276 | -48.18 |
| 1D0DA | 60 | 95 | -240.33 |
| 1D2ZC | 108 | 229 | -78.88 |
| 1D3BG | 73 | 134 | 14.54 |
| 1D3BL | 90 | 151 | -34.88 |
| 1DBFC | 127 | 251 | -47.01 |
| 1DCSA | 279 | 511 | 16.03 |
| 1DEVD | 39 | 66 | 182.41 |
| 1DG6A | 149 | 249 | -213.67 |
| 1DGWA | 178 | 309 | -57.85 |
| 1DIXA | 208 | 376 | -262.31 |
| 1DPJB | 29 | 66 | -28.22 |
| 1DSZB | 84 | 140 | -254.54 |
| 1DTDB | 61 | 92 | -225.13 |
| 1DY2A | 168 | 320 | -52.5 |
| 1EAID | 61 | 102 | -156.76 |
| 1EB6A | 177 | 361 | -253.92 |
| 1EDNA | 21 | 36 | -211.79 |

| | | | |
|---|---|---|---|
| 1EEXB | 178 | 368 | 216.96 |
| 1EFND | 104 | 218 | 43.85 |
| 1EGIB | 143 | 262 | -256.56 |
| 1EGWC | 72 | 131 | -79.87 |
| 1EJFB | 110 | 189 | -287.16 |
| 1EODA | 100 | 201 | 80.04 |
| 1EZJA | 114 | 259 | -230.53 |
| 1F1UB | 321 | 620 | 137.41 |
| 1F2LD | 70 | 133 | -231.11 |
| 1F2TA | 145 | 244 | -2.07 |
| 1F32A | 127 | 246 | -38.2 |
| 1F7CA | 182 | 369 | -63.91 |
| 1F94A | 63 | 105 | -498.3 |
| 1FD3C | 40 | 68 | -122.6 |
| 1FO0B | 112 | 189 | -118.7 |
| 1FQJC | 38 | 71 | 85.69 |
| 1FR3A | 67 | 105 | 55.93 |
| 1FS1B | 116 | 257 | -41.86 |
| 1FS1C | 41 | 87 | 6.41 |
| 1FSGC | 233 | 427 | -42.15 |
| 1FV1C | 20 | 20 | 82.73 |
| 1FVIA | 264 | 505 | 267.67 |
| 1G2CR | 37 | 81 | -21.16 |
| 1G2CU | 50 | 118 | 8.5 |
| 1G2RA | 94 | 183 | -57.46 |
| 1G31G | 107 | 185 | 206.02 |
| 1G4IA | 123 | 277 | -733.7 |
| 1G8EA | 98 | 199 | -150.45 |
| 1GA6A | 369 | 768 | 559.09 |
| 1GBGA | 214 | 367 | 154.56 |
| 1GEFD | 109 | 210 | 34.18 |
| 1GHQB | 129 | 201 | -37.12 |
| 1GL2D | 55 | 130 | -109.46 |
| 1GOUA | 109 | 190 | 97.54 |
| 1GPPA | 217 | 364 | 57.94 |
| 1GUQA | 347 | 691 | -92.57 |
| 1GVDA | 52 | 95 | -102.91 |
| 1GWEA | 498 | 943 | 247.54 |
| 1GY7A | 121 | 217 | -106.95 |
| 1GYXA | 76 | 140 | -37.41 |
| 1H2KS | 24 | 35 | -12.71 |
| 1H6FA | 184 | 323 | -29.6 |
| 1H7ZA | 191 | 309 | 157.78 |
| 1H8PB | 88 | 155 | -233.41 |
| 1H97A | 147 | 316 | -27.06 |
| 1HDKA | 139 | 230 | 27.98 |
| 1HH5A | 68 | 116 | -122.52 |

| | | | |
|---|---|---|---|
| 1HK0X | 173 | 308 | -288.8 |
| 1HWGC | 191 | 331 | -189.71 |
| 1I27A | 73 | 135 | 21.22 |
| 1I8FD | 73 | 127 | 7.43 |
| 1ID0A | 146 | 275 | 158.43 |
| 1IEJA | 329 | 618 | -373.13 |
| 1IGQA | 54 | 85 | 29.29 |
| 1IK9C | 28 | 46 | -15.51 |
| 1IQ5B | 24 | 47 | 49.07 |
| 1IQZA | 81 | 141 | 166.82 |
| 1IWMB | 177 | 328 | 54.27 |
| 1IX9B | 205 | 410 | -29.44 |
| 1IXHA | 321 | 639 | 358.2 |
| 1J0PA | 108 | 195 | -359.66 |
| 1J34B | 123 | 225 | -335.49 |
| 1J3AA | 129 | 247 | 156.86 |
| 1JB0C | 80 | 140 | -172.59 |
| 1JB0E | 69 | 112 | 127.26 |
| 1JB0I | 38 | 75 | 121.12 |
| 1JB0L | 151 | 314 | 414.01 |
| 1JB0M | 31 | 72 | 24.46 |
| 1JB0X | 29 | 72 | -26.29 |
| 1JDHB | 38 | 61 | -86.94 |
| 1JEKA | 40 | 102 | -12.33 |
| 1JEKB | 34 | 76 | -167.11 |
| 1JF8A | 130 | 235 | 33.04 |
| 1JFBA | 399 | 808 | 146.66 |
| 1JM1A | 202 | 365 | 160.19 |
| 1JMTA | 98 | 176 | -91.61 |
| 1JMTB | 23 | 37 | 105.67 |
| 1JNXX | 207 | 431 | 38.98 |
| 1JO0A | 97 | 180 | -43.41 |
| 1JR8B | 105 | 212 | -206.58 |
| 1JWQA | 179 | 391 | -53.02 |
| 1JY2O | 51 | 88 | -15.88 |
| 1JY2P | 44 | 78 | -114.46 |
| 1K61A | 60 | 127 | -82.45 |
| 1KNGA | 144 | 266 | 292.73 |
| 1KNQB | 171 | 363 | -124.53 |
| 1KTGB | 132 | 242 | -100.3 |
| 1KVEA | 63 | 101 | 12.49 |
| 1KWFA | 363 | 753 | -46.72 |
| 1KYFA | 247 | 449 | -90.76 |
| 1L2HA | 144 | 246 | -98.29 |
| 1L2WI | 57 | 83 | 60.51 |
| 1L3AA | 166 | 286 | 292.62 |
| 1L9LA | 74 | 145 | -195.83 |

| | | | |
|---|---|---|---|
| 1LGHA | 56 | 121 | 66.54 |
| 1LGHB | 43 | 92 | 39.6 |
| 1LJ2C | 22 | 42 | 15.66 |
| 1LKKA | 105 | 182 | -7.98 |
| 1LN0A | 92 | 188 | -151.01 |
| 1LO7A | 140 | 264 | 95.51 |
| 1LQTB | 454 | 894 | 686 |
| 1LRWD | 83 | 156 | 98.2 |
| 1LTAC | 45 | 85 | -59.92 |
| 1M0DD | 129 | 234 | 24.93 |
| 1M15A | 356 | 741 | -19.79 |
| 1M1QA | 90 | 182 | -213.66 |
| 1M2DB | 101 | 189 | 172.22 |
| 1M2ZE | 21 | 25 | -33.61 |
| 1M40A | 263 | 657 | 62.51 |
| 1M45B | 25 | 53 | -61.06 |
| 1M46B | 25 | 55 | -110.55 |
| 1M4ZB | 203 | 387 | -59.19 |
| 1M8AB | 61 | 106 | -160.05 |
| 1M93A | 46 | 87 | 88.16 |
| 1MCTI | 28 | 46 | -201.05 |
| 1MF7A | 194 | 422 | -34.87 |
| 1MHHF | 62 | 120 | 83.5 |
| 1MJ4A | 80 | 163 | 111.44 |
| 1MJ5A | 298 | 609 | 277.69 |
| 1MJCA | 69 | 116 | 94.94 |
| 1MN8A | 95 | 193 | 151.63 |
| 1MTPB | 35 | 46 | 69.59 |
| 1MVFE | 44 | 62 | 51.56 |
| 1MXEF | 25 | 49 | 1.12 |
| 1MZWB | 31 | 63 | 3.11 |
| 1N0WB | 33 | 53 | -37.05 |
| 1N13A | 46 | 63 | 101 |
| 1N1JA | 87 | 176 | -116.06 |
| 1N7SD | 66 | 148 | -85.52 |
| 1NBUC | 118 | 218 | 108.01 |
| 1NBWD | 113 | 233 | 133.09 |
| 1NEZG | 122 | 191 | -76.56 |
| 1NFPA | 228 | 461 | -169.57 |
| 1NO5B | 102 | 185 | -27.73 |
| 1NOFA | 383 | 760 | 541.65 |
| 1NPIA | 61 | 105 | -283.95 |
| 1NRJB | 191 | 341 | 15.55 |
| 1NS4B | 346 | 668 | -50.68 |
| 1NT3A | 108 | 176 | -270.11 |
| 1NVPC | 47 | 77 | -46.39 |
| 1NWAA | 168 | 302 | -1.12 |

| | | | |
|---|---|---|---|
| 1NYCA | 111 | 198 | -198.5 |
| 1O06A | 20 | 41 | -28.79 |
| 1O7IB | 114 | 183 | 128.61 |
| 1O7JA | 325 | 628 | 430.48 |
| 1OAIA | 59 | 116 | -13.23 |
| 1OAUI | 22 | 15 | 6.65 |
| 1OB8B | 127 | 243 | -76.76 |
| 1OC0B | 37 | 67 | -391.3 |
| 1OC7A | 364 | 739 | 223.82 |
| 1ONKA | 249 | 522 | 86.07 |
| 1OOHB | 126 | 275 | -263.41 |
| 1OQJA | 90 | 171 | -22.47 |
| 1OR7C | 66 | 143 | -30.19 |
| 1ORYB | 40 | 90 | 2.24 |
| 1OV9B | 48 | 103 | -92.53 |
| 1P27D | 92 | 158 | 99.33 |
| 1P28B | 117 | 243 | -197.11 |
| 1P57A | 110 | 205 | -81.1 |
| 1P5ZB | 229 | 444 | -239.13 |
| 1P6OA | 156 | 332 | 1.4 |
| 1PJMA | 20 | 21 | 47.27 |
| 1PJNA | 21 | 26 | -8.73 |
| 1PK4A | 79 | 133 | -160.68 |
| 1PM4A | 117 | 196 | -72.46 |
| 1PPJJ | 33 | 64 | -33.47 |
| 1PSRA | 100 | 213 | -235.65 |
| 1PWGA | 345 | 717 | 214.8 |
| 1Q0AA | 91 | 182 | -220.6 |
| 1Q35A | 317 | 646 | -30.21 |
| 1Q5YD | 80 | 153 | -194.54 |
| 1QDDA | 144 | 234 | -145.01 |
| 1QGRB | 24 | 55 | -71.12 |
| 1QLWB | 317 | 610 | 432.43 |
| 1QOAB | 98 | 183 | 69.48 |
| 1QQP2 | 216 | 364 | 134.37 |
| 1QSMA | 150 | 279 | -76.32 |
| 1QTXB | 20 | 42 | -11.03 |
| 1QVYA | 138 | 228 | 58.45 |
| 1QWYA | 234 | 423 | 119.41 |
| 1R2QA | 170 | 345 | -69 |
| 1R44A | 202 | 403 | -21.97 |
| 1R6JA | 82 | 154 | 17.72 |
| 1R77A | 99 | 174 | 119.16 |
| 1R8HA | 87 | 149 | -40.46 |
| 1R8SA | 160 | 296 | -160.28 |
| 1RG8B | 141 | 246 | -91.49 |
| 1RH6A | 55 | 94 | -37.39 |

| | | | |
|---|---|---|---|
| 1RKIB | 97 | 170 | -245.75 |
| 1RLKA | 116 | 223 | -51.22 |
| 1RMDA | 116 | 212 | -257.64 |
| 1RQLB | 257 | 558 | 267.15 |
| 1RSSA | 140 | 267 | -41.24 |
| 1RTQA | 291 | 594 | -16.87 |
| 1RUTX | 157 | 245 | -153.68 |
| 1RWEA | 21 | 40 | -165.58 |
| 1RYLA | 157 | 317 | -160.89 |
| 1RYQA | 64 | 104 | -214.96 |
| 1S0PB | 176 | 396 | 9.84 |
| 1S3MB | 165 | 335 | -292.32 |
| 1S4CA | 150 | 267 | 19.6 |
| 1S5NA | 386 | 797 | 434.26 |
| 1S6CB | 21 | 37 | 49.14 |
| 1S7KA | 158 | 304 | -71.01 |
| 1SAUA | 114 | 229 | -112.82 |
| 1SCZA | 233 | 440 | 235.82 |
| 1SFSA | 213 | 432 | 59.37 |
| 1SHUX | 181 | 374 | -12.5 |
| 1SHXB | 138 | 238 | -26.07 |
| 1SJWA | 142 | 273 | 201.01 |
| 1SLUA | 131 | 194 | -15.23 |
| 1SN4A | 64 | 113 | -320.38 |
| 1SPPB | 112 | 192 | -89.49 |
| 1SVFC | 62 | 150 | 58.37 |
| 1SVFD | 35 | 77 | -31.93 |
| 1SYYA | 317 | 706 | -116.03 |
| 1SZ7A | 159 | 303 | -28.76 |
| 1T3YA | 131 | 254 | -136.1 |
| 1T61C | 222 | 406 | -136.83 |
| 1T6OA | 49 | 94 | -61.01 |
| 1T8KA | 77 | 155 | 65.72 |
| 1T8ZD | 47 | 104 | -265.63 |
| 1TAFB | 70 | 140 | -14.7 |
| 1TAGA | 314 | 655 | -471.11 |
| 1TB3A | 332 | 678 | -143.07 |
| 1TG0A | 66 | 117 | 88.07 |
| 1TJLD | 145 | 332 | -370.99 |
| 1TTWB | 20 | 32 | -8.97 |
| 1TU3I | 45 | 99 | -161.73 |
| 1TUKA | 67 | 142 | -343.93 |
| 1TVDB | 116 | 199 | -2.09 |
| 1TVGA | 136 | 236 | -27.39 |
| 1TWFI | 122 | 195 | -340.19 |
| 1U07A | 90 | 145 | 179.33 |
| 1U2HA | 96 | 162 | 24.85 |

| | | | |
|---|---|---|---|
| 1U2WB | 107 | 211 | -119.57 |
| 1U36A | 100 | 165 | -54.91 |
| 1U4GA | 298 | 618 | 250.45 |
| 1U5DD | 108 | 186 | -188.88 |
| 1U6HB | 24 | 50 | -33.78 |
| 1UFBD | 127 | 286 | 62.87 |
| 1UFIB | 46 | 93 | 7.96 |
| 1UG4A | 60 | 101 | -321.61 |
| 1ULIB | 177 | 317 | -64.37 |
| 1UNNC | 111 | 212 | -129.83 |
| 1UNRA | 113 | 191 | -180.85 |
| 1UOYA | 64 | 120 | -291.52 |
| 1URRA | 97 | 169 | -28.09 |
| 1US0A | 314 | 657 | -87.29 |
| 1USDA | 42 | 92 | -113.22 |
| 1UVQC | 20 | 19 | 44.42 |
| 1UWCB | 261 | 498 | -96.74 |
| 1V2ZA | 106 | 216 | -107.02 |
| 1V6PB | 62 | 99 | -457.77 |
| 1V74A | 107 | 212 | -93.79 |
| 1VBWA | 68 | 121 | -32.06 |
| 1VF6C | 51 | 116 | -105.71 |
| 1VKEE | 119 | 262 | -129.41 |
| 1VKIB | 166 | 320 | 62.49 |
| 1VLSA | 146 | 337 | -183.79 |
| 1VMBA | 107 | 211 | -210.08 |
| 1VPPX | 20 | 25 | -52.52 |
| 1VZIB | 125 | 204 | -151.52 |
| 1W0NA | 120 | 206 | 157.72 |
| 1W1DA | 144 | 263 | -115.48 |
| 1W5QA | 321 | 647 | 331.02 |
| 1W66A | 218 | 408 | 209.19 |
| 1WKWB | 20 | 32 | -37.2 |
| 1WLZA | 85 | 174 | -109.98 |
| 1WM3A | 72 | 125 | 46.65 |
| 1WMIC | 88 | 174 | 100.78 |
| 1WP7C | 61 | 158 | -145.81 |
| 1WQJB | 80 | 126 | -316.04 |
| 1WU4A | 374 | 766 | 117.28 |
| 1WWBX | 103 | 164 | -41.04 |
| 1X0TA | 106 | 204 | -152.96 |
| 1X6IB | 87 | 179 | -53.27 |
| 1X8QA | 184 | 340 | -170.77 |
| 1XAKA | 68 | 113 | -186.87 |
| 1XBIA | 118 | 231 | 107.07 |
| 1XBWB | 96 | 174 | -21.17 |
| 1XD7A | 116 | 271 | 34.95 |

| | | | |
|---|---|---|---|
| 1XG5D | 257 | 535 | 54.85 |
| 1XJUA | 156 | 311 | -295.66 |
| 1XLQC | 106 | 187 | -25.91 |
| 1XU1A | 137 | 219 | -7.44 |
| 1XVXA | 311 | 614 | 295.9 |
| 1Y66C | 44 | 92 | -205.47 |
| 1Y93A | 158 | 318 | 76.48 |
| 1Y96C | 86 | 143 | -16.46 |
| 1YD2A | 90 | 152 | 20.38 |
| 1YDIB | 24 | 52 | -49.25 |
| 1YF6M | 301 | 621 | 299.09 |
| 1YK0E | 42 | 87 | -121.49 |
| 1YK4A | 52 | 89 | 56.38 |
| 1YSQA | 181 | 336 | 11.52 |
| 1YYPB | 20 | 30 | -40.45 |
| 1Z1SA | 136 | 258 | 26.14 |
| 1Z2UA | 150 | 278 | 98.94 |
| 1Z3EB | 67 | 146 | -42.43 |
| 1Z5GA | 208 | 408 | 160.68 |
| 1Z8IA | 39 | 77 | -16.54 |
| 1Z96B | 35 | 77 | 51.9 |
| 1ZAVZ | 27 | 62 | -32.1 |
| 1ZBA4 | 49 | 63 | -57.22 |
| 1ZBXB | 120 | 216 | -80.55 |
| 1ZMMC | 31 | 48 | -282.16 |
| 1ZMPD | 25 | 46 | -249 |
| 1ZMQD | 32 | 53 | -312.01 |
| 1ZOQC | 47 | 84 | -26.51 |
| 1ZPSB | 128 | 232 | -117.69 |
| 1ZT3A | 80 | 137 | -181.41 |
| 1ZUUA | 56 | 95 | 18.99 |
| 1ZV8C | 46 | 97 | -102.01 |
| 1ZVBB | 33 | 68 | -46.09 |
| 1ZVZB | 22 | 38 | -19.2 |
| 1ZW0B | 59 | 126 | -110.48 |
| 1ZW2B | 21 | 36 | -20.26 |
| 1ZZKA | 80 | 152 | 39.68 |
| 2A06I | 42 | 55 | 88.69 |
| 2A2FX | 296 | 659 | -253.4 |
| 2A40C | 22 | 36 | -17.25 |
| 2A4DA | 139 | 254 | 200.76 |
| 2A6ZA | 222 | 396 | 36.63 |
| 2A83A | 276 | 503 | -331.03 |
| 2A9IA | 105 | 215 | 32.59 |
| 2ADVB | 28 | 32 | 17.09 |
| 2AF0A | 146 | 292 | -78.47 |
| 2AHYA | 104 | 254 | 0.91 |

| | | | |
|---|---|---|---|
| 2AKFB | 32 | 66 | -78.43 |
| 2ANXB | 146 | 287 | -138.95 |
| 2AO9H | 114 | 206 | -119.71 |
| 2ASKB | 101 | 164 | -76.38 |
| 2AYDA | 76 | 114 | 20.88 |
| 2B3GA | 117 | 209 | 83.95 |
| 2B3GB | 24 | 45 | 94.55 |
| 2B5AA | 77 | 162 | 0.53 |
| 2B5ID | 122 | 180 | -298.95 |
| 2BAYC | 55 | 97 | 20.02 |
| 2BCMA | 138 | 231 | 16.32 |
| 2BCXB | 27 | 59 | -58.78 |
| 2BE6F | 22 | 41 | -37.77 |
| 2BEZF | 42 | 62 | -4.35 |
| 2BF6A | 449 | 820 | 292 |
| 2BJNA | 143 | 278 | -63.41 |
| 2BKRA | 212 | 411 | -227.51 |
| 2BL8B | 82 | 176 | -40.78 |
| 2BLFB | 81 | 132 | 86.54 |
| 2BMJA | 174 | 319 | -200.46 |
| 2BO1A | 101 | 197 | 148.67 |
| 2BOSB | 68 | 128 | -96.14 |
| 2BOXA | 135 | 234 | -543.36 |
| 2BT9A | 90 | 132 | 14.54 |
| 2BZVA | 148 | 255 | 67.41 |
| 2BZWB | 27 | 57 | 24.26 |
| 2C4FT | 75 | 129 | -12.36 |
| 2C5KP | 22 | 40 | -52.36 |
| 2C5KT | 89 | 176 | -226.71 |
| 2C5LC | 97 | 172 | -26.92 |
| 2CAKA | 154 | 258 | 365.41 |
| 2CARB | 194 | 361 | 6.93 |
| 2CB5B | 453 | 925 | -207.83 |
| 2CB8A | 86 | 172 | -81.07 |
| 2CCVA | 99 | 168 | -51.4 |
| 2CG7A | 90 | 158 | -288.06 |
| 2CHHA | 113 | 202 | 120.97 |
| 2CIUA | 123 | 210 | -32.43 |
| 2CJTC | 127 | 212 | -57.72 |
| 2CKLA | 98 | 178 | -95.33 |
| 2CNQA | 301 | 649 | 110.28 |
| 2CO4A | 123 | 210 | 76.07 |
| 2CPGC | 42 | 76 | -34.52 |
| 2CS7A | 55 | 92 | 9.76 |
| 2CVIB | 83 | 146 | -165.71 |
| 2CWSA | 227 | 436 | 274.16 |
| 2CZQB | 205 | 414 | 185.09 |

| | | | |
|---|---|---|---|
| 2D10H | 20 | 29 | -23.88 |
| 2D1KC | 29 | 40 | 46.12 |
| 2D48A | 129 | 263 | -577.07 |
| 2D7ED | 103 | 171 | 229.15 |
| 2D80A | 318 | 650 | 201.14 |
| 2DJFA | 118 | 203 | -139.04 |
| 2DKOA | 146 | 274 | -163.4 |
| 2DKOB | 103 | 174 | 38.95 |
| 2DS5B | 44 | 76 | -186.2 |
| 2DYOB | 36 | 69 | -78.23 |
| 2E4TA | 509 | 1048 | 380.02 |
| 2EBBA | 96 | 191 | -110.85 |
| 2EGIH | 97 | 177 | -79.84 |
| 2ENDA | 137 | 278 | 58.49 |
| 2EQ7C | 37 | 69 | 74.76 |
| 2ERLA | 40 | 77 | -244.68 |
| 2EULA | 156 | 326 | 1.82 |
| 2EV0B | 131 | 277 | -73.52 |
| 2EY4D | 75 | 131 | 195.04 |
| 2F31B | 20 | 38 | 7.59 |
| 2F3NC | 62 | 141 | -86.3 |
| 2F5YB | 82 | 137 | -2.76 |
| 2F69A | 244 | 432 | 334.13 |
| 2FCWB | 78 | 163 | -396.36 |
| 2FGOA | 81 | 140 | -67.64 |
| 2FHZA | 106 | 201 | 56.96 |
| 2FOTC | 23 | 51 | 31.96 |
| 2FQMC | 67 | 113 | -52.69 |
| 2FR5A | 136 | 262 | 9.49 |
| 2FU5B | 107 | 173 | -96.96 |
| 2FYZB | 32 | 73 | -31.74 |
| 2G0CA | 68 | 133 | 92.35 |
| 2G9ZA | 297 | 577 | -157.21 |
| 2GB4A | 231 | 449 | -198.23 |
| 2GE7B | 108 | 186 | 57.02 |
| 2GGCA | 263 | 508 | -36.36 |
| 2GJ3A | 119 | 233 | 110.08 |
| 2GKGA | 122 | 264 | 43.92 |
| 2GKRI | 51 | 97 | -204.81 |
| 2GMYE | 145 | 308 | -77.59 |
| 2GOMA | 61 | 133 | -135.32 |
| 2GPED | 48 | 82 | -16.23 |
| 2GRRB | 157 | 314 | 29.78 |
| 2GZGA | 83 | 158 | -25.76 |
| 2GZVA | 91 | 156 | 132.45 |
| 2H3LB | 98 | 160 | 112.73 |
| 2H5CA | 198 | 387 | 68.92 |

| | | | |
|---|---|---|---|
| 2H62C | 84 | 139 | -461.67 |
| 2H9DA | 84 | 182 | 36.08 |
| 2H9EC | 52 | 86 | -228.84 |
| 2HEWF | 128 | 212 | -121.62 |
| 2HEYR | 138 | 218 | -474.62 |
| 2HLQA | 100 | 182 | -519.78 |
| 2HNTE | 67 | 102 | -33.07 |
| 2HPJA | 99 | 201 | 3.6 |
| 2HQHF | 22 | 33 | -19.99 |
| 2HQLC | 91 | 162 | -42.65 |
| 2HS1B | 99 | 159 | 175.4 |
| 2HT0B | 93 | 181 | 28.33 |
| 2HVWA | 147 | 268 | -40.81 |
| 2HWNB | 45 | 73 | 60.59 |
| 2HWNF | 20 | 39 | -43.25 |
| 2I15C | 115 | 246 | -175.32 |
| 2I2FA | 262 | 465 | 224.2 |
| 2I2QA | 122 | 223 | 124.62 |
| 2I52F | 119 | 208 | 105.05 |
| 2I53A | 254 | 556 | -187.67 |
| 2I8TA | 149 | 253 | 109.85 |
| 2IDQA | 105 | 183 | 100.28 |
| 2IHCC | 117 | 204 | -65.11 |
| 2IHSD | 20 | 31 | -65.43 |
| 2IJKA | 57 | 126 | -202.11 |
| 2IQJA | 118 | 219 | -110.94 |
| 2IUMA | 211 | 371 | 340.48 |
| 2IYBF | 63 | 98 | -184.74 |
| 2IYJB | 75 | 128 | 78.36 |
| 2J1KB | 113 | 171 | 35.35 |
| 2J3WC | 135 | 254 | 61.82 |
| 2J4WD | 34 | 48 | -217.37 |
| 2J6YA | 85 | 188 | 76.86 |
| 2J73A | 103 | 171 | -0.84 |
| 2J97A | 98 | 161 | 131.07 |
| 2J9UB | 47 | 71 | 38.28 |
| 2J9WA | 101 | 211 | -108.81 |
| 2JCQA | 150 | 273 | -131.19 |
| 2JDAB | 142 | 249 | 41.05 |
| 2JDCA | 145 | 266 | 47.07 |
| 2JFRA | 234 | 444 | 185.01 |
| 2JGSC | 112 | 184 | -8.48 |
| 2JJUB | 112 | 184 | 75.25 |
| 2MLTB | 26 | 53 | 4.2 |
| 2NL9B | 23 | 48 | 18.68 |
| 2NLRA | 222 | 409 | 292.55 |
| 2NMLA | 100 | 206 | -70.13 |

| | | | |
|---|---|---|---|
| 2NNUB | 20 | 31 | -5.63 |
| 2NPSD | 63 | 160 | -68.15 |
| 2NPTC | 103 | 174 | 109.34 |
| 2NQ2B | 300 | 666 | 90.73 |
| 2NQDA | 109 | 177 | 134.18 |
| 2NSZA | 129 | 273 | -157.93 |
| 2NUDD | 23 | 28 | -11.2 |
| 2NVAB | 369 | 701 | 164.5 |
| 2NXVA | 249 | 461 | -219.3 |
| 2O31A | 67 | 117 | 130.59 |
| 2O60B | 20 | 42 | -9.53 |
| 2O6PB | 119 | 212 | -26.62 |
| 2O8MC | 23 | 22 | 29.17 |
| 2ODBB | 35 | 48 | 57.63 |
| 2OFCA | 141 | 260 | -96.85 |
| 2OFKB | 183 | 385 | -227.5 |
| 2OKQA | 127 | 229 | 73.64 |
| 2OKRF | 24 | 43 | -14.47 |
| 2ON5A | 206 | 411 | 91.33 |
| 2OPEC | 111 | 218 | -58.25 |
| 2ORWB | 173 | 324 | 6.98 |
| 2OUGA | 141 | 295 | 45.78 |
| 2OVCA | 30 | 57 | 14.7 |
| 2OY3A | 98 | 183 | -285.45 |
| 2P04B | 105 | 179 | 157.94 |
| 2P0BA | 142 | 288 | -270.88 |
| 2P28B | 214 | 344 | -890.99 |
| 2P39A | 142 | 243 | 128.24 |
| 2P4FA | 193 | 382 | -247.03 |
| 2P84A | 133 | 237 | -59.09 |
| 2PBDV | 28 | 34 | 111.24 |
| 2PJPA | 121 | 220 | -273.47 |
| 2PK8A | 94 | 176 | 25.89 |
| 2PL6A | 71 | 119 | -236.59 |
| 2PLXB | 26 | 53 | -267.12 |
| 2POSD | 94 | 188 | -186.9 |
| 2PQ3A | 74 | 156 | 112.58 |
| 2PQMA | 338 | 677 | 332.47 |
| 2PQNB | 20 | 39 | 31.61 |
| 2PQRD | 45 | 75 | 67.85 |
| 2PRGC | 32 | 57 | -130.68 |
| 2PSMB | 117 | 237 | -255.63 |
| 2PU3A | 207 | 410 | -367.23 |
| 2PV2A | 103 | 190 | 75.35 |
| 2PVBA | 107 | 233 | -68.76 |
| 2PY2E | 127 | 250 | -386.78 |
| 2Q6QB | 65 | 149 | -310.43 |

| | | | |
|---|---|---|---|
| 2QCPX | 80 | 136 | 26.96 |
| 2QF4B | 167 | 288 | -23.26 |
| 2QFAC | 45 | 91 | -87.73 |
| 2QIFB | 68 | 125 | -23.88 |
| 2QIYB | 133 | 241 | -90.33 |
| 2QIYD | 25 | 42 | 29.63 |
| 2QKHB | 32 | 69 | -38.99 |
| 2QMEI | 24 | 32 | 58.06 |
| 2QPWA | 147 | 232 | 213.3 |
| 2QSKA | 95 | 182 | -252.91 |
| 2QTVD | 36 | 50 | 12.23 |
| 2QUXK | 122 | 191 | 24.85 |
| 2QVKA | 121 | 196 | 98.86 |
| 2R25B | 133 | 249 | 35.04 |
| 2R2VD | 33 | 72 | -107.56 |
| 2R31A | 236 | 505 | 309.45 |
| 2RBLA | 89 | 133 | 13.11 |
| 2RH2A | 58 | 103 | 90.27 |
| 2RK5A | 86 | 152 | -39.35 |
| 2RKLB | 50 | 111 | -49.02 |
| 2RKYB | 21 | 22 | -30.13 |
| 2RKZP | 21 | 23 | -11.8 |
| 2UTGB | 70 | 158 | 71.45 |
| 2UU8A | 237 | 430 | 273.84 |
| 2UUYB | 52 | 78 | -208.59 |
| 2UV4A | 144 | 269 | 68.91 |
| 2UVPD | 176 | 344 | -210.91 |
| 2UWJF | 32 | 63 | -15.79 |
| 2UWRA | 79 | 156 | -580.59 |
| 2UX9E | 67 | 111 | -6.07 |
| 2V33B | 91 | 154 | -163.27 |
| 2V3GA | 273 | 557 | 145.73 |
| 2V52M | 30 | 57 | -36.87 |
| 2V6YB | 72 | 143 | -18.17 |
| 2V7QI | 47 | 79 | -8.84 |
| 2V85A | 74 | 118 | -120.34 |
| 2V8BA | 279 | 581 | 265.02 |
| 2V9LA | 274 | 566 | 200.18 |
| 2V9VA | 135 | 273 | -158.41 |
| 2VB1A | 129 | 282 | -335.18 |
| 2VC8A | 72 | 131 | 137.07 |
| 2VE8G | 61 | 114 | 94.16 |
| 2VFXH | 203 | 353 | -146.55 |
| 2VGOC | 41 | 66 | 56.92 |
| 2VJFC | 64 | 100 | -24.5 |
| 2VQPA | 255 | 462 | 71.59 |
| 2VT1A | 21 | 30 | -19.3 |

| | | | |
|---|---|---|---|
| 2VXNA | 249 | 508 | 22.88 |
| 2VXTI | 156 | 269 | -60.03 |
| 2VZCA | 127 | 270 | -23.41 |
| 2W1RA | 117 | 221 | -18.91 |
| 2W39A | 298 | 570 | 59.98 |
| 2W3GA | 150 | 266 | 222.34 |
| 2W3NA | 230 | 436 | 238.93 |
| 2W50B | 97 | 193 | -253.87 |
| 2W6AB | 56 | 124 | -132.55 |
| 2W72C | 141 | 311 | 179.49 |
| 2W83C | 67 | 136 | -84.6 |
| 2W8XA | 72 | 144 | -647.84 |
| 2W91A | 636 | 1221 | 32.02 |
| 2WASA | 113 | 222 | 80.19 |
| 2WAXD | 28 | 51 | 31.59 |
| 2WDQD | 105 | 224 | 19.01 |
| 2WF7A | 218 | 427 | 225.38 |
| 2WFUA | 22 | 42 | -146.68 |
| 2WFVB | 23 | 34 | 55.69 |
| 2WJRA | 204 | 373 | -95.91 |
| 2WLUA | 166 | 336 | -0.78 |
| 2WLVA | 144 | 296 | 153.35 |
| 2WV6D | 92 | 154 | -113.39 |
| 2WX0C | 31 | 46 | -83.92 |
| 2WX3B | 42 | 76 | -84.84 |
| 2WY7Q | 66 | 141 | -158.53 |
| 2WYTA | 153 | 280 | -79.74 |
| 2WZ1A | 197 | 345 | 50.68 |
| 2WZ8A | 135 | 240 | 69.06 |
| 2X04A | 77 | 162 | 76.57 |
| 2X2OA | 117 | 226 | 74.35 |
| 2X32A | 174 | 320 | 173.66 |
| 2X5YA | 171 | 306 | -135.68 |
| 2X6PB | 29 | 63 | -106.69 |
| 2X9CA | 62 | 135 | 2.2 |
| 2X9GD | 249 | 492 | 205.84 |
| 2XFRA | 487 | 965 | 484.53 |
| 2XHFB | 154 | 312 | 163.6 |
| 2XJPA | 258 | 481 | 221.66 |
| 2XMJB | 63 | 120 | -70.74 |
| 2XOLB | 172 | 356 | -90.74 |
| 2XPNB | 20 | 35 | -11.3 |
| 2XPPB | 24 | 46 | -1.25 |
| 2XQQB | 89 | 157 | -129.81 |
| 2XS2A | 88 | 143 | 157.4 |
| 2XTCB | 62 | 119 | 61.75 |
| 2XTSB | 204 | 349 | 305.72 |

| | | | |
|---|---|---|---|
| 2XTSC | 389 | 709 | 352.18 |
| 2XTVA | 178 | 366 | 120.59 |
| 2XVOC | 180 | 302 | -11.91 |
| 2XWVA | 309 | 660 | 67.12 |
| 2XXNA | 143 | 240 | -136.08 |
| 2XZER | 20 | 39 | -26.26 |
| 2Y1BA | 86 | 167 | -35.68 |
| 2Y1KA | 525 | 1078 | 19.25 |
| 2Y20D | 54 | 103 | -0.61 |
| 2Y2YA | 95 | 143 | -65 |
| 2Y3NB | 60 | 122 | 24.95 |
| 2Y5PD | 72 | 112 | 106.21 |
| 2Y71A | 138 | 273 | -20.56 |
| 2Y78A | 122 | 216 | 267.9 |
| 2Y7BA | 134 | 237 | -271.54 |
| 2Y8FB | 112 | 201 | -207.16 |
| 2Y9UA | 67 | 148 | -130.97 |
| 2Y9WC | 136 | 219 | 93.6 |
| 2YA3B | 68 | 88 | 65.1 |
| 2YALA | 39 | 84 | -57.54 |
| 2YBFB | 21 | 33 | -23.41 |
| 2YF2D | 46 | 83 | -23.85 |
| 2YH5A | 121 | 235 | 111.37 |
| 2YLEB | 21 | 31 | 33.87 |
| 2Z0WA | 72 | 109 | 176.61 |
| 2Z14A | 116 | 193 | 85.06 |
| 2Z1CC | 71 | 115 | 106.5 |
| 2Z34D | 22 | 25 | 10.45 |
| 2Z3QB | 81 | 128 | 9.16 |
| 2Z4UA | 261 | 504 | -123.3 |
| 2Z57B | 65 | 102 | 135.39 |
| 2Z5BB | 132 | 231 | -22.32 |
| 2Z7FI | 50 | 80 | -265.23 |
| 2ZA4B | 89 | 187 | -100.55 |
| 2ZAEC | 105 | 183 | 107.86 |
| 2ZFDB | 116 | 203 | -34.93 |
| 2ZK9X | 185 | 359 | -265.29 |
| 2ZNRA | 178 | 327 | -225.93 |
| 2ZS0D | 145 | 308 | 106.4 |
| 2ZSIB | 60 | 117 | 2.32 |
| 2ZU0D | 42 | 80 | 40.07 |
| 2ZVMC | 249 | 424 | -16.29 |
| 2ZYZC | 96 | 169 | -28.01 |
| 2ZZDA | 119 | 197 | 63.75 |
| 2ZZDK | 152 | 293 | -31.77 |
| 3A1FA | 163 | 334 | 37.21 |
| 3A4UA | 228 | 393 | 70.07 |

| | | | |
|---|---|---|---|
| 3A57A | 154 | 272 | -23.45 |
| 3A8GB | 212 | 386 | 471.53 |
| 3A9FA | 78 | 159 | 95.88 |
| 3AA1C | 21 | 29 | 42.99 |
| 3AA6C | 23 | 31 | 71.36 |
| 3ABDY | 22 | 33 | 40.51 |
| 3ADLA | 76 | 130 | 52.8 |
| 3ADYA | 102 | 196 | 130.11 |
| 3AIAA | 200 | 386 | -247.38 |
| 3AJ4B | 112 | 206 | -226.2 |
| 3AJ6A | 283 | 553 | -570.59 |
| 3AJBB | 26 | 39 | 5.9 |
| 3AJIB | 73 | 158 | -26.86 |
| 3ALRD | 55 | 97 | -38.22 |
| 3ASLA | 68 | 110 | -187.8 |
| 3AU4B | 32 | 69 | -10.48 |
| 3AUBA | 58 | 93 | 10.81 |
| 3AWUA | 278 | 567 | -47.19 |
| 3AX2G | 66 | 138 | -37.44 |
| 3B0DB | 95 | 208 | 79.21 |
| 3B0FA | 43 | 97 | 9.47 |
| 3B0ZA | 41 | 82 | -4.03 |
| 3B1JD | 23 | 38 | -36.94 |
| 3B4UB | 288 | 578 | 218.75 |
| 3BBBF | 151 | 282 | 0.48 |
| 3BC1B | 52 | 110 | -56.46 |
| 3BD1B | 63 | 132 | 60.95 |
| 3BFXB | 248 | 486 | -104.22 |
| 3BRVA | 39 | 76 | -101.82 |
| 3BS2A | 146 | 248 | -255.48 |
| 3BT4A | 85 | 138 | -364.44 |
| 3BTPB | 28 | 47 | -8.53 |
| 3BWZA | 171 | 289 | 189.82 |
| 3BYAB | 20 | 40 | -0.27 |
| 3BYPA | 82 | 147 | 106 |
| 3BZVA | 24 | 31 | -9.79 |
| 3C0TB | 23 | 47 | -24.5 |
| 3C4MD | 20 | 40 | -64.33 |
| 3C4SA | 57 | 94 | 9.99 |
| 3C5TB | 25 | 53 | -36.66 |
| 3C6AA | 198 | 378 | -129.38 |
| 3C6WC | 52 | 79 | -128.9 |
| 3C70A | 256 | 549 | -143.4 |
| 3C7TA | 259 | 464 | 244.93 |
| 3C8IA | 127 | 223 | 241.89 |
| 3C8YA | 574 | 1188 | -277.26 |
| 3C9AC | 48 | 73 | -107.74 |

| | | | |
|---|---|---|---|
| 3CAAB | 32 | 38 | 38.2 |
| 3CC0A | 106 | 179 | 105.91 |
| 3CI9B | 45 | 111 | -119.84 |
| 3CJID | 35 | 45 | 28.66 |
| 3CJSB | 72 | 126 | 214.9 |
| 3CNKA | 89 | 146 | 81.39 |
| 3CP5A | 116 | 212 | -19.7 |
| 3CSQD | 326 | 634 | 51.22 |
| 3CT6B | 128 | 252 | 37.32 |
| 3CTRA | 75 | 132 | 117.48 |
| 3CU0B | 247 | 465 | 251.99 |
| 3CX5I | 57 | 105 | -33.71 |
| 3CZ1B | 115 | 231 | -252.34 |
| 3D06A | 179 | 302 | -77.86 |
| 3D2QD | 69 | 110 | -82.63 |
| 3D32A | 118 | 216 | 110.83 |
| 3D55B | 84 | 167 | 7.01 |
| 3D5YA | 314 | 548 | 315.17 |
| 3D9NB | 140 | 314 | -20.05 |
| 3DBOA | 34 | 57 | 34.31 |
| 3DDCB | 133 | 224 | -21.72 |
| 3DDTC | 44 | 73 | -101.67 |
| 3DHAA | 254 | 488 | 73.5 |
| 3DJ9A | 107 | 179 | 66.05 |
| 3DK9A | 462 | 898 | 225.52 |
| 3DS4A | 77 | 147 | 52.27 |
| 3DVEB | 23 | 47 | 9.1 |
| 3DVUC | 24 | 58 | 14.59 |
| 3DXEB | 27 | 42 | 9.24 |
| 3E1EF | 141 | 264 | 168.45 |
| 3E21A | 40 | 82 | -38.6 |
| 3E4HA | 29 | 48 | -169.65 |
| 3E5AB | 33 | 50 | -10.23 |
| 3E5TA | 228 | 427 | 277.11 |
| 3E7HB | 101 | 175 | 85.51 |
| 3E8MA | 164 | 331 | 82.05 |
| 3E8YX | 30 | 52 | -231.21 |
| 3E9TB | 113 | 191 | 161.37 |
| 3EA6A | 215 | 425 | -108.26 |
| 3ECHC | 24 | 44 | -56.99 |
| 3EGGD | 66 | 90 | 113.52 |
| 3EIPA | 84 | 151 | 43.6 |
| 3EJ3C | 64 | 114 | 100.29 |
| 3ELSA | 145 | 252 | -15.14 |
| 3EO5A | 169 | 306 | 370.16 |
| 3EOPB | 161 | 329 | -51.59 |
| 3ERXA | 123 | 211 | 90.48 |

| | | | |
|---|---|---|---|
| 3ESID | 124 | 217 | 27.3 |
| 3EX7I | 62 | 99 | -19.46 |
| 3EYIA | 64 | 125 | -85.63 |
| 3F1IC | 24 | 49 | -1.7 |
| 3F1PA | 114 | 212 | -18 |
| 3F4TA | 200 | 390 | -264.22 |
| 3F6YA | 235 | 444 | -419.34 |
| 3FBLA | 82 | 166 | -31.81 |
| 3FGHA | 67 | 133 | -79.17 |
| 3FJUB | 65 | 124 | -606.66 |
| 3FO3B | 519 | 1057 | -591.85 |
| 3FPUA | 100 | 157 | -355.1 |
| 3FS3A | 197 | 412 | -192.32 |
| 3FSAA | 122 | 234 | -35.7 |
| 3FWBB | 54 | 115 | -88.74 |
| 3FX7A | 84 | 178 | -254.75 |
| 3FXDA | 49 | 104 | -57.16 |
| 3G21A | 77 | 165 | 236.45 |
| 3G3BG | 71 | 106 | -104.07 |
| 3G43F | 45 | 104 | -48.35 |
| 3G46A | 146 | 334 | -13.51 |
| 3G48A | 110 | 177 | 144.96 |
| 3G4ZC | 117 | 259 | -41.14 |
| 3G66A | 190 | 330 | 106.84 |
| 3G7CA | 109 | 244 | -270.06 |
| 3G7ZC | 34 | 60 | -52.67 |
| 3G80A | 73 | 157 | -22.18 |
| 3G91A | 260 | 513 | -128.06 |
| 3G9WC | 39 | 71 | -52.01 |
| 3G9YA | 29 | 52 | -74.63 |
| 3GA3A | 133 | 242 | -382.44 |
| 3GA4A | 156 | 277 | -50.7 |
| 3GA8A | 67 | 111 | -29.34 |
| 3GAEB | 253 | 539 | 83.76 |
| 3GB3A | 229 | 391 | 161.35 |
| 3GCMD | 21 | 22 | 31.85 |
| 3GE3B | 304 | 680 | -423.32 |
| 3GJ8B | 27 | 43 | -94.23 |
| 3GLAA | 97 | 167 | 125.54 |
| 3GP2B | 20 | 41 | -7.62 |
| 3GP6A | 155 | 274 | 98.36 |
| 3GS2B | 30 | 40 | -13.77 |
| 3GWNB | 113 | 248 | -136.19 |
| 3GXVD | 26 | 56 | -35.4 |
| 3H0TC | 22 | 33 | -463.6 |
| 3H0WB | 61 | 90 | -50.24 |
| 3H3GB | 22 | 38 | -64.73 |

| | | | |
|---|---|---|---|
| 3H5JA | 168 | 318 | 193.03 |
| 3H63A | 315 | 653 | -206.47 |
| 3H7HB | 95 | 165 | 47.67 |
| 3H7XD | 54 | 113 | -91.92 |
| 3H87B | 138 | 268 | 28.02 |
| 3H8AE | 28 | 40 | 72.82 |
| 3H8DG | 38 | 62 | -12.03 |
| 3HCTA | 104 | 182 | -183.35 |
| 3HF5D | 109 | 206 | -35.9 |
| 3HFEB | 26 | 56 | -60.49 |
| 3HFNB | 60 | 102 | -84.19 |
| 3HIAA | 66 | 105 | 37.35 |
| 3HIHB | 216 | 411 | -236.37 |
| 3HILB | 62 | 131 | 4.77 |
| 3HMRA | 90 | 165 | -303.87 |
| 3HNXA | 108 | 199 | -65.43 |
| 3HPWB | 101 | 179 | 76.04 |
| 3HRZC | 362 | 596 | -562.61 |
| 3HTKA | 60 | 151 | -239.02 |
| 3HTKC | 254 | 500 | -129.71 |
| 3HTUF | 32 | 63 | -140.22 |
| 3I94A | 243 | 477 | -19.39 |
| 3IBMB | 146 | 239 | -48.43 |
| 3IEZB | 95 | 186 | -76.06 |
| 3IGMA | 56 | 94 | 11.5 |
| 3IKKB | 125 | 215 | 136.15 |
| 3ILYB | 139 | 266 | 9.78 |
| 3IMOB | 105 | 185 | 35.69 |
| 3IOLB | 26 | 51 | 0.19 |
| 3IP0A | 158 | 294 | 106.47 |
| 3ITMA | 307 | 691 | -506.6 |
| 3IVVA | 140 | 243 | -111.65 |
| 3IWIA | 354 | 653 | 591.62 |
| 3IXSB | 31 | 43 | -40.78 |
| 3IXSE | 101 | 180 | -9.98 |
| 3JQHA | 23 | 47 | -64.32 |
| 3JUDA | 145 | 253 | 226.23 |
| 3K43B | 145 | 272 | -243.54 |
| 3K67A | 156 | 283 | 160.72 |
| 3K6TC | 54 | 111 | -21.95 |
| 3K80C | 103 | 161 | 1.3 |
| 3KCQB | 204 | 373 | -88.3 |
| 3KEOB | 207 | 396 | 130.79 |
| 3KF6A | 136 | 244 | 11.92 |
| 3KFAA | 136 | 244 | 11.92 |
| 3KGRC | 99 | 160 | 114.04 |
| 3KIHD | 91 | 152 | 35.61 |

| | | | |
|---|---|---|---|
| 3KM3B | 156 | 267 | 130.31 |
| 3KM5A | 179 | 316 | 165.46 |
| 3KT9A | 102 | 170 | -48.52 |
| 3KU3B | 172 | 319 | -244.26 |
| 3KUPC | 59 | 104 | -25.79 |
| 3KUVA | 132 | 248 | 259.11 |
| 3KZ5A | 47 | 80 | -30.57 |
| 3L1EA | 105 | 169 | 49.49 |
| 3L2CA | 85 | 161 | -40.43 |
| 3L46A | 90 | 165 | 13.02 |
| 3L4QC | 163 | 378 | -563.09 |
| 3L4RA | 151 | 263 | -127.39 |
| 3L9QA | 171 | 335 | -16.3 |
| 3LBLC | 91 | 183 | -14.12 |
| 3LCNC | 25 | 48 | -119.88 |
| 3LE1B | 85 | 176 | -13.67 |
| 3LE4A | 55 | 76 | 140.35 |
| 3LJDA | 126 | 227 | -136.85 |
| 3LK4C | 29 | 40 | 49.23 |
| 3LMSB | 74 | 135 | -441.48 |
| 3LOEA | 30 | 46 | -240.46 |
| 3LOIA | 161 | 256 | 101.2 |
| 3LR2A | 125 | 268 | 142.9 |
| 3LU9C | 25 | 30 | 15.37 |
| 3LUCA | 135 | 256 | 19.85 |
| 3LW3B | 137 | 232 | 45.9 |
| 3LW6A | 241 | 456 | -331.8 |
| 3LYEA | 285 | 562 | 260.37 |
| 3M20A | 59 | 111 | 85.97 |
| 3M21B | 66 | 129 | 0.55 |
| 3M4WH | 31 | 46 | 22.81 |
| 3M5QA | 357 | 689 | 154.4 |
| 3M7FA | 107 | 191 | -202.03 |
| 3M7OA | 137 | 221 | -188.05 |
| 3M8JB | 88 | 179 | -6.54 |
| 3M91C | 45 | 92 | -111.9 |
| 3M91D | 31 | 57 | -50.16 |
| 3M9QB | 87 | 158 | -113.26 |
| 3MA2B | 117 | 203 | -75.21 |
| 3MABB | 88 | 174 | -70.72 |
| 3MAZA | 99 | 175 | 109.71 |
| 3MCBB | 58 | 96 | 20.34 |
| 3MD1A | 83 | 145 | 45.96 |
| 3MD7A | 270 | 497 | 125.64 |
| 3MEAA | 162 | 249 | 191.84 |
| 3MEZA | 111 | 187 | -36.89 |
| 3MHPC | 26 | 32 | 89.39 |

| | | | |
|---|---|---|---|
| 3MHSB | 91 | 189 | -221.59 |
| 3MJHB | 34 | 57 | 16.9 |
| 3MJKX | 121 | 207 | -262.62 |
| 3ML1B | 109 | 169 | -7.46 |
| 3MLHB | 213 | 361 | -24.75 |
| 3MLSS | 20 | 33 | -25.86 |
| 3MMHA | 167 | 317 | -69.27 |
| 3MMYF | 51 | 87 | -9.14 |
| 3MN5S | 20 | 34 | -12.72 |
| 3MN7S | 20 | 31 | -6.31 |
| 3MQIC | 89 | 134 | 121.86 |
| 3MTSB | 61 | 104 | -131.23 |
| 3MU7A | 273 | 537 | 185.81 |
| 3MUDD | 48 | 99 | -106.07 |
| 3MVCB | 154 | 331 | -147.83 |
| 3MXNA | 151 | 272 | -47.77 |
| 3N1FC | 94 | 159 | 29.83 |
| 3N3FA | 54 | 87 | 30.14 |
| 3N4WB | 81 | 153 | 64.99 |
| 3N7DB | 55 | 90 | 12.8 |
| 3N7SC | 91 | 178 | -243.92 |
| 3NA0D | 28 | 40 | -41.26 |
| 3NDDB | 35 | 48 | 84.06 |
| 3NFCA | 94 | 151 | 157.54 |
| 3NFLC | 89 | 149 | 63.36 |
| 3NGGA | 46 | 79 | -253.61 |
| 3NJ0A | 177 | 321 | 69.3 |
| 3NK4C | 22 | 25 | -110.25 |
| 3NPVA | 249 | 510 | 125.44 |
| 3NR5A | 157 | 285 | -138.27 |
| 3NSWG | 94 | 152 | -202.19 |
| 3NV0A | 196 | 376 | 71.16 |
| 3NVSA | 426 | 859 | 293.93 |
| 3NZ3A | 100 | 154 | -29.28 |
| 3O1FB | 81 | 152 | -41.62 |
| 3O2RA | 144 | 319 | -154.04 |
| 3O3KA | 90 | 184 | -84.36 |
| 3O4PA | 314 | 575 | 269.1 |
| 3O6QD | 44 | 65 | 47.47 |
| 3O7AA | 52 | 88 | -174.86 |
| 3O8VA | 102 | 169 | 45.24 |
| 3OA2D | 298 | 606 | 147.8 |
| 3OADD | 171 | 285 | 3.43 |
| 3OAKC | 31 | 66 | 37.04 |
| 3OBLB | 132 | 228 | 128.11 |
| 3OBQA | 141 | 247 | 249.09 |
| 3OGNB | 124 | 239 | -309.4 |

| | | | |
|---|---|---|---|
| 3OIQB | 30 | 57 | -10 |
| 3OJ3O | 32 | 68 | -119.57 |
| 3ONHA | 113 | 206 | -23.6 |
| 3ONRJ | 68 | 113 | -10.92 |
| 3OP8B | 85 | 145 | -269.98 |
| 3OPKB | 115 | 195 | -47.77 |
| 3OQ2B | 102 | 184 | -2.51 |
| 3OR7C | 103 | 240 | 28.42 |
| 3OSKB | 123 | 199 | 62.32 |
| 3OSVC | 132 | 221 | 158.41 |
| 3OWTC | 20 | 32 | -15.35 |
| 3P57P | 105 | 232 | -640.58 |
| 3P8CE | 64 | 128 | -114.88 |
| 3PB6X | 313 | 613 | 177.93 |
| 3PC6B | 103 | 186 | -59.17 |
| 3PD7B | 94 | 184 | -98.85 |
| 3PGZB | 114 | 193 | -182.61 |
| 3PH0C | 53 | 120 | -175.74 |
| 3PHTA | 115 | 206 | -198.12 |
| 3PISA | 39 | 69 | -101.48 |
| 3PIVA | 156 | 346 | -250.83 |
| 3PLVC | 21 | 36 | 21.45 |
| 3PLXA | 25 | 26 | 15.48 |
| 3PM2A | 173 | 316 | -390.63 |
| 3PMCB | 132 | 280 | -344.98 |
| 3PMEA | 415 | 751 | -303.88 |
| 3PMTA | 55 | 89 | 20.16 |
| 3PP2A | 112 | 196 | 4.76 |
| 3PRPC | 161 | 306 | -3.73 |
| 3PT8B | 152 | 347 | -110.72 |
| 3PUCA | 98 | 170 | 86.53 |
| 3Q2BA | 120 | 232 | -136.39 |
| 3Q46A | 178 | 342 | 162.71 |
| 3Q4OA | 169 | 380 | -140.5 |
| 3Q7TB | 100 | 175 | -50.58 |
| 3QKSC | 21 | 45 | -18.76 |
| 3QL9A | 125 | 252 | -540.29 |
| 3QPIA | 173 | 310 | -54.9 |
| 3QQQB | 149 | 306 | 129.98 |
| 3QR7B | 115 | 145 | 174.53 |
| 3QV1H | 20 | 33 | -78.39 |
| 3QWGB | 90 | 176 | 40.99 |
| 3QZBA | 132 | 221 | -44.04 |
| 3QZLA | 121 | 215 | -69.08 |
| 3R1WA | 180 | 371 | 247.36 |
| 3R2DA | 132 | 266 | -19.92 |
| 3R42A | 156 | 275 | 285.76 |

| | | | |
|---|---|---|---|
| 3R62A | 125 | 197 | -249.41 |
| 3R85E | 20 | 35 | -19.94 |
| 3R85F | 21 | 46 | -92.37 |
| 3R87A | 132 | 249 | -81.36 |
| 3RAYA | 159 | 253 | -27.29 |
| 3RF3B | 258 | 544 | 0.74 |
| 3RHGA | 363 | 765 | -81.35 |
| 3RI8A | 182 | 326 | 101.07 |
| 3RJPA | 96 | 174 | -24.37 |
| 3RKQA | 58 | 115 | -76.25 |
| 3RLGA | 278 | 572 | 66.12 |
| 3RLOA | 196 | 342 | -60.52 |
| 3RLSA | 152 | 241 | 205.17 |
| 3RMIA | 105 | 226 | -204.91 |
| 3RO2B | 22 | 27 | -23.85 |
| 3RO3A | 159 | 353 | -186.48 |
| 3RPEA | 197 | 384 | 190.61 |
| 3RPPC | 216 | 429 | 70.79 |
| 3RQ9A | 78 | 153 | 21.59 |
| 3RT3C | 94 | 212 | 62.4 |
| 3RTLA | 114 | 201 | -21 |
| 3RWNA | 155 | 266 | 54.85 |
| 3RY2A | 121 | 210 | 104.56 |
| 3RZNA | 206 | 439 | 118.78 |
| 3S0AA | 119 | 252 | -298.41 |
| 3S6PE | 33 | 72 | 25.66 |
| 3S90D | 26 | 52 | -24.12 |
| 3SGRC | 24 | 29 | 42.92 |
| 3SH4A | 195 | 330 | 245.23 |
| 3SJMB | 55 | 109 | 41.44 |
| 3SK2B | 132 | 217 | 43.45 |
| 3SL9H | 20 | 39 | -114.92 |
| 3SMVA | 233 | 474 | -6.4 |
| 3SO6A | 137 | 250 | -58.1 |
| 3SOVA | 306 | 569 | -121.96 |
| 3SQFB | 94 | 168 | 5.13 |
| 3SSBD | 34 | 47 | -68.21 |
| 3SSBJ | 29 | 41 | -55.82 |
| 3SSRB | 99 | 185 | 189.75 |
| 3STDC | 162 | 295 | -36.69 |
| 3SWFA | 56 | 118 | -110.27 |
| 3SZSA | 42 | 74 | -352.84 |
| 3T0HA | 209 | 410 | -185.6 |
| 3T2CA | 389 | 803 | 687.28 |
| 3T5XB | 30 | 54 | -58.05 |
| 3T7LA | 74 | 129 | -303.05 |
| 3T7YA | 94 | 173 | 207.44 |

| | | | |
|---|---|---|---|
| 3THMF | 77 | 134 | -504.61 |
| 3TJ5B | 27 | 48 | 12.49 |
| 3TOTB | 202 | 383 | -61.63 |
| 3TQ2A | 35 | 79 | -73.41 |
| 3TQ5A | 113 | 186 | 274.07 |
| 3TRSA | 37 | 40 | 65.76 |
| 3TUOB | 103 | 171 | -11.3 |
| 3TVJI | 35 | 58 | -269.36 |
| 3TVKA | 172 | 313 | 19.65 |
| 3TVQA | 152 | 287 | 9.52 |
| 3TWEB | 25 | 51 | -109.38 |
| 3TZ1B | 24 | 43 | -19.24 |
| 3U1JA | 20 | 22 | -27.53 |
| 3U23A | 56 | 92 | -31.35 |
| 3U3PA | 164 | 271 | -455.97 |
| 3U5WA | 126 | 209 | -117.1 |
| 3U65A | 314 | 655 | -132.68 |
| 3U7CA | 258 | 495 | 115.16 |
| 3U7QA | 477 | 974 | 130.51 |
| 3U8EA | 216 | 396 | 178.27 |
| 3U99A | 123 | 233 | 34.47 |
| 3U9DD | 21 | 36 | -34.03 |
| 3U9QA | 257 | 532 | -145.05 |
| 3U9ZC | 20 | 32 | -49.13 |
| 3UANB | 254 | 499 | -84.54 |
| 3UAWA | 233 | 519 | 113.67 |
| 3UEJA | 65 | 101 | -174.21 |
| 3UK3D | 54 | 91 | -186.4 |
| 3UKWC | 22 | 27 | 34.12 |
| 3UL0C | 23 | 26 | -20.78 |
| 3UL1A | 21 | 22 | 11.02 |
| 3ULBB | 97 | 157 | -54.84 |
| 3UTMC | 32 | 35 | 138.35 |
| 3V0DB | 325 | 634 | -56.73 |
| 3V1EB | 43 | 83 | -60.71 |
| 3V2IA | 187 | 363 | 108.47 |
| 3V3WA | 397 | 772 | 333.96 |
| 3V46A | 160 | 297 | -26.1 |
| 3V4HA | 131 | 225 | -105.47 |
| 3V4YB | 41 | 59 | 23.82 |
| 3VBHB | 245 | 421 | 181.75 |
| 3VC8B | 85 | 161 | -87.88 |
| 3VCAA | 397 | 737 | 65.26 |
| 3VEJA | 40 | 82 | -32.75 |
| 3VH5W | 75 | 167 | -23.33 |
| 3VHSB | 26 | 42 | -20.1 |
| 3VIIA | 472 | 976 | 310.1 |

| | | | |
|---|---|---|---|
| 3VK6A | 96 | 148 | -103.27 |
| 3VM9B | 153 | 329 | 52.05 |
| 3VMTB | 218 | 486 | -255.65 |
| 3VMXC | 48 | 106 | -197.04 |
| 3VP9B | 77 | 173 | -265.98 |
| 3VQJA | 214 | 413 | 232.71 |
| 3VTGA | 200 | 387 | -158.71 |
| 3VTQF | 37 | 74 | -211.64 |
| 3VU9A | 215 | 398 | -102.22 |
| 3VUPA | 351 | 726 | -50.67 |
| 3VUXF | 32 | 55 | -153.78 |
| 3VV4A | 153 | 288 | -185.75 |
| 3VVID | 21 | 44 | -15.77 |
| 3VVVA | 108 | 173 | 55.03 |
| 3VWXB | 218 | 447 | 87.22 |
| 3VYXA | 95 | 170 | -104.21 |
| 3VZ9B | 103 | 186 | -167.88 |
| 3VZAC | 62 | 113 | 21.99 |
| 3VZAF | 31 | 50 | 36.31 |
| 3W06A | 267 | 556 | 195.36 |
| 3W07A | 215 | 468 | 292.37 |
| 3W15A | 336 | 626 | -201.56 |
| 3W19C | 38 | 78 | -123.93 |
| 3W42B | 199 | 384 | -150.02 |
| 3W5HA | 272 | 499 | 341.77 |
| 3WDNA | 125 | 239 | 65.93 |
| 3WE7C | 267 | 531 | 77.15 |
| 3WGDF | 109 | 200 | -210.25 |
| 3WH1A | 203 | 403 | 62.99 |
| 3WH2A | 132 | 250 | -312.17 |
| 3WHJA | 111 | 229 | -89.47 |
| 3WMIA | 57 | 129 | -80.43 |
| 3WMQA | 94 | 150 | -113.48 |
| 3WN7M | 24 | 47 | -20.97 |
| 3WOUA | 207 | 372 | -275.96 |
| 3WTTA | 119 | 194 | 65.37 |
| 3WU2Y | 27 | 55 | 77.99 |
| 3WUPA | 31 | 56 | -0.87 |
| 3WUTK | 43 | 89 | -150.85 |
| 3WWQF | 39 | 65 | -65.62 |
| 3WXBA | 269 | 539 | 99.11 |
| 3WY9D | 26 | 58 | -74.83 |
| 3WYGD | 20 | 32 | -28.02 |
| 3X0IA | 159 | 281 | 193.47 |
| 3X0TA | 112 | 192 | 20.97 |
| 3X2MA | 180 | 340 | -166.07 |
| 3X2WS | 20 | 29 | 1.22 |

| | | | |
|---|---|---|---|
| 3X34A | 87 | 184 | -55.84 |
| 3ZBOB | 235 | 505 | -201.77 |
| 3ZD2B | 119 | 176 | -200.42 |
| 3ZDFC | 21 | 33 | -76.18 |
| 3ZDLB | 26 | 59 | 23.5 |
| 3ZG8A | 39 | 69 | -13.35 |
| 3ZIAJ | 36 | 57 | -5.9 |
| 3ZIIA | 82 | 139 | -23.13 |
| 3ZITB | 78 | 139 | -53.64 |
| 3ZOQB | 48 | 81 | 8.15 |
| 3ZQ6D | 307 | 624 | 58.11 |
| 3ZQSA | 186 | 337 | 110.33 |
| 3ZS9C | 31 | 68 | -95.1 |
| 3ZSJA | 138 | 232 | -47.75 |
| 3ZUCA | 153 | 273 | 40.84 |
| 3ZWLF | 39 | 78 | -142.99 |
| 3ZXCA | 71 | 115 | -210.35 |
| 3ZZLA | 70 | 112 | -37.47 |
| 3ZZOA | 93 | 163 | -457.8 |
| 4A02A | 166 | 310 | 192.47 |
| 4A1SE | 28 | 40 | 36.44 |
| 4A29A | 247 | 525 | -52.33 |
| 4A69C | 69 | 128 | -76.73 |
| 4A6HA | 108 | 173 | -20.35 |
| 4A8UA | 159 | 291 | 143.78 |
| 4A8XB | 27 | 42 | 44.01 |
| 4AANA | 323 | 646 | 691.59 |
| 4ABLA | 181 | 366 | -74.64 |
| 4ACJA | 167 | 306 | 120.42 |
| 4ADNB | 207 | 391 | -111.35 |
| 4AEQA | 90 | 150 | 7.66 |
| 4AFFA | 110 | 188 | 38.45 |
| 4AJXH | 29 | 52 | -35.95 |
| 4ALAC | 78 | 107 | 39.15 |
| 4ANNA | 176 | 354 | -63.49 |
| 4AORE | 34 | 64 | -123.38 |
| 4AS8A | 228 | 422 | 101.17 |
| 4AUVF | 23 | 45 | -97.63 |
| 4AUVG | 27 | 53 | -118.24 |
| 4AYAA | 59 | 112 | -34.72 |
| 4AYOA | 434 | 887 | 384.52 |
| 4B0MA | 131 | 216 | 88.82 |
| 4B1UM | 30 | 53 | -63.39 |
| 4B1YM | 30 | 57 | -99.9 |
| 4B5OA | 192 | 359 | -70.93 |
| 4B6DB | 57 | 87 | -105.29 |
| 4B6IC | 102 | 214 | -161.24 |

| | | | |
|---|---|---|---|
| 4B9GA | 146 | 240 | 185.59 |
| 4BB2B | 33 | 43 | 17.62 |
| 4BBKA | 121 | 212 | -34.1 |
| 4BD8D | 136 | 270 | -33.52 |
| 4BFHA | 30 | 49 | -219.99 |
| 4BFOA | 106 | 174 | 58.72 |
| 4BHXA | 90 | 184 | -48.2 |
| 4BI8A | 94 | 182 | -112.76 |
| 4BJ0A | 167 | 293 | 218.95 |
| 4BJJB | 85 | 114 | 5.99 |
| 4BJSD | 49 | 95 | 19.34 |
| 4BJTA | 152 | 296 | -13.94 |
| 4BL0E | 45 | 69 | 76.48 |
| 4BPZA | 251 | 460 | 44.82 |
| 4BRCB | 360 | 709 | -9.96 |
| 4BWQF | 25 | 43 | 65.28 |
| 4C0FD | 100 | 173 | 14.51 |
| 4C5GB | 30 | 42 | 0.08 |
| 4C9SB | 262 | 474 | 286.54 |
| 4CAYC | 23 | 30 | 9.11 |
| 4CC2A | 64 | 112 | -2.33 |
| 4CCGX | 75 | 129 | -91.94 |
| 4CE7A | 345 | 707 | 35.75 |
| 4CFQC | 85 | 183 | -80.71 |
| 4CFQQ | 27 | 54 | 21.69 |
| 4CHIA | 320 | 658 | 455.43 |
| 4CJ0A | 534 | 1063 | 286.31 |
| 4CJ2C | 62 | 102 | -68.67 |
| 4CK4A | 160 | 320 | -194.95 |
| 4CN0B | 93 | 141 | 52.09 |
| 4CNGA | 156 | 306 | 79.08 |
| 4CNNB | 247 | 493 | 165.72 |
| 4CO6E | 39 | 73 | -28.64 |
| 4CPAI | 37 | 66 | -208.68 |
| 4CRUB | 270 | 585 | -279.55 |
| 4CRWB | 158 | 327 | -302.03 |
| 4CU4B | 21 | 28 | 91.47 |
| 4CXFA | 154 | 311 | 77.5 |
| 4CXFB | 29 | 53 | -24.65 |
| 4CXIA | 130 | 246 | 40.4 |
| 4CYDF | 21 | 35 | -43.66 |
| 4CZ7D | 31 | 50 | -71.44 |
| 4D0GC | 46 | 99 | -61.97 |
| 4D0KB | 105 | 190 | -49.63 |
| 4D0QA | 161 | 277 | -145.12 |
| 4D2HG | 35 | 56 | -121.5 |
| 4D63A | 138 | 229 | 60.35 |

| | | | |
|---|---|---|---|
| 4D6KB | 69 | 149 | -128.69 |
| 4D7ZA | 28 | 30 | 26.2 |
| 4D8BA | 254 | 490 | 236.71 |
| 4DACD | 26 | 56 | -22.2 |
| 4DEYB | 29 | 65 | -119.64 |
| 4DHXD | 69 | 142 | -239.75 |
| 4DJ9B | 22 | 42 | 0.36 |
| 4DKAD | 82 | 122 | 34.79 |
| 4DKKA | 106 | 201 | -12.93 |
| 4DKNB | 214 | 384 | 266.8 |
| 4DNCD | 42 | 69 | -43.03 |
| 4DR8B | 187 | 332 | -43 |
| 4DS7G | 36 | 65 | -95.08 |
| 4DT4A | 160 | 272 | 245.05 |
| 4DW5A | 158 | 294 | -365.73 |
| 4DXRB | 25 | 32 | 24.56 |
| 4E17B | 28 | 51 | -88.24 |
| 4E3YB | 240 | 461 | 164.05 |
| 4E4SF | 114 | 195 | -48.91 |
| 4E6KI | 56 | 106 | -155.83 |
| 4E74A | 101 | 169 | 19.92 |
| 4EFOA | 89 | 159 | 25.37 |
| 4EGUB | 115 | 204 | -27.36 |
| 4EHQG | 20 | 29 | -21.53 |
| 4EICA | 93 | 183 | 51.53 |
| 4EO1A | 67 | 107 | 21.96 |
| 4ES7A | 166 | 291 | 6.23 |
| 4ESPA | 130 | 253 | 185.46 |
| 4ETNA | 148 | 302 | -34.64 |
| 4EVYA | 147 | 275 | -133.94 |
| 4EX6A | 219 | 435 | 703.03 |
| 4EZFB | 66 | 101 | -20.48 |
| 4F0ZC | 30 | 35 | -12.73 |
| 4F42A | 84 | 168 | -83.21 |
| 4F6MA | 116 | 185 | -200.53 |
| 4F7UE | 78 | 142 | -84.44 |
| 4F8LA | 138 | 238 | 180.46 |
| 4FASF | 49 | 82 | -74.42 |
| 4FBWC | 21 | 22 | -2.14 |
| 4FCJB | 127 | 230 | -6.64 |
| 4FDDB | 20 | 32 | -15.79 |
| 4FI5A | 71 | 153 | -171.26 |
| 4FK5E | 91 | 146 | 20.69 |
| 4FP5H | 98 | 174 | 17.29 |
| 4FQGA | 186 | 403 | -585.44 |
| 4FTXA | 158 | 288 | 39.2 |
| 4FU6A | 95 | 166 | 185.5 |

| | | | |
|---|---|---|---|
| 4FVDA | 138 | 238 | 6.2 |
| 4FXXC | 91 | 165 | 117.87 |
| 4G0YA | 147 | 274 | -86.49 |
| 4G28B | 87 | 185 | -193.18 |
| 4G2EA | 151 | 282 | 200.68 |
| 4G3OA | 53 | 115 | -7.18 |
| 4G4KB | 100 | 180 | -203.88 |
| 4G6FF | 27 | 57 | -89.73 |
| 4G78A | 147 | 324 | -317.82 |
| 4G9SB | 111 | 188 | 122.82 |
| 4GA2A | 144 | 335 | -154.35 |
| 4GCOA | 120 | 275 | -209.39 |
| 4GF3B | 28 | 53 | -42.51 |
| 4GFTA | 63 | 120 | -60.73 |
| 4GGFS | 89 | 184 | -90.35 |
| 4GHOB | 96 | 159 | 129.45 |
| 4GJZA | 228 | 415 | 120.03 |
| 4GLSE | 95 | 157 | -194.43 |
| 4GR6A | 104 | 227 | -23.28 |
| 4GRNA | 116 | 217 | 133.78 |
| 4GT8A | 133 | 238 | -80.3 |
| 4GUSC | 20 | 25 | 19.14 |
| 4GV5C | 42 | 79 | -231.48 |
| 4GVBA | 77 | 154 | -459.81 |
| 4GVIA | 333 | 656 | 140.09 |
| 4GXZC | 174 | 357 | 40.54 |
| 4H10A | 60 | 136 | -101.69 |
| 4H25C | 22 | 22 | 34.01 |
| 4H2VC | 26 | 42 | 57.5 |
| 4H2WC | 78 | 160 | 128.1 |
| 4H6CB | 177 | 317 | 219.79 |
| 4H7RD | 31 | 75 | 12.66 |
| 4H7WA | 187 | 361 | -140.24 |
| 4HASB | 103 | 201 | -34.04 |
| 4HE6A | 89 | 144 | 121.44 |
| 4HI8A | 170 | 331 | -45.7 |
| 4HI9B | 72 | 115 | -67.28 |
| 4HJLB | 192 | 338 | -401.44 |
| 4HJPA | 281 | 534 | -143.12 |
| 4HKGA | 80 | 163 | 75.92 |
| 4HLSB | 105 | 190 | -135.68 |
| 4HLYA | 106 | 189 | 16.09 |
| 4HMSA | 202 | 354 | 112.87 |
| 4HNOA | 285 | 607 | -284.04 |
| 4HOIC | 110 | 208 | -144.9 |
| 4HPMD | 82 | 148 | -155.87 |
| 4HR6A | 41 | 67 | 40.64 |

| | | | |
|---|---|---|---|
| 4HS1A | 84 | 149 | -43.37 |
| 4HTMA | 28 | 52 | -62.26 |
| 4HVKA | 369 | 760 | 258.44 |
| 4HVUA | 57 | 93 | 37.42 |
| 4HW5A | 77 | 159 | -312.72 |
| 4HWFA | 87 | 187 | 3.04 |
| 4HY4B | 95 | 171 | -5.38 |
| 4I0XD | 83 | 179 | -141.35 |
| 4I4SD | 146 | 251 | 81.1 |
| 4I8HA | 223 | 413 | -279.8 |
| 4ICVA | 84 | 148 | 2.14 |
| 4ID8A | 65 | 114 | 30.8 |
| 4IF6B | 26 | 38 | 66.34 |
| 4IGIA | 197 | 413 | 384.75 |
| 4IN0A | 141 | 262 | 43.44 |
| 4INKA | 203 | 377 | 229.08 |
| 4IQMA | 313 | 610 | -13.26 |
| 4IRGA | 98 | 188 | -86.8 |
| 4IUMA | 127 | 232 | 86.63 |
| 4IZXA | 139 | 244 | 93.64 |
| 4J2CA | 108 | 233 | -41.36 |
| 4J2KB | 167 | 275 | 19.84 |
| 4J32B | 101 | 204 | -182.58 |
| 4J4AI | 22 | 44 | -41.01 |
| 4J4AL | 21 | 44 | -6.79 |
| 4J5OB | 109 | 205 | 42.85 |
| 4J5RB | 139 | 279 | -153.48 |
| 4J7BC | 49 | 74 | 23 |
| 4JB3A | 218 | 449 | 107.73 |
| 4JEAD | 106 | 195 | -255.8 |
| 4JG2A | 185 | 318 | 137.84 |
| 4JIFA | 130 | 223 | 24.86 |
| 4JJRA | 284 | 538 | 30.34 |
| 4JK8A | 220 | 410 | -48.7 |
| 4JO6Z | 25 | 48 | -6.31 |
| 4JPNA | 75 | 170 | -117.44 |
| 4JPRA | 76 | 170 | -163.56 |
| 4JQUB | 54 | 105 | -144.42 |
| 4JS0B | 29 | 38 | 192.43 |
| 4JVOA | 336 | 651 | 140.94 |
| 4JZZA | 336 | 592 | -624.16 |
| 4K02B | 127 | 224 | 94.02 |
| 4K12B | 82 | 177 | -99.32 |
| 4K2PD | 232 | 444 | -180.53 |
| 4K45A | 102 | 190 | -52.17 |
| 4K5BD | 144 | 287 | -16.02 |
| 4K94C | 195 | 309 | 83.11 |

| | | | |
|---|---|---|---|
| 4KDIA | 157 | 285 | 21.98 |
| 4KDIC | 65 | 110 | -26.88 |
| 4KFSA | 202 | 390 | 114.05 |
| 4KHBC | 106 | 169 | 79.64 |
| 4KNQA | 122 | 220 | -40.12 |
| 4KQDD | 115 | 201 | -159.65 |
| 4KQPA | 230 | 447 | 241.31 |
| 4KRUA | 214 | 417 | -190.17 |
| 4KU0D | 96 | 169 | 61.29 |
| 4KXLC | 134 | 259 | 64.6 |
| 4KXVA | 620 | 1258 | 4.8 |
| 4L0KC | 204 | 405 | -292.57 |
| 4L7XA | 58 | 97 | -198.52 |
| 4L82B | 167 | 285 | -129.34 |
| 4L8IA | 114 | 241 | -350.78 |
| 4L9DA | 82 | 137 | 5.69 |
| 4L9UA | 49 | 101 | -113.8 |
| 4LD6A | 117 | 212 | 93.9 |
| 4LGJA | 256 | 528 | 15.28 |
| 4LH9A | 40 | 66 | 30.8 |
| 4LJ0A | 65 | 98 | -69.1 |
| 4LKSA | 157 | 277 | 2.26 |
| 4LKUD | 25 | 48 | -54.95 |
| 4LMSC | 68 | 109 | -112.6 |
| 4LOQL | 22 | 21 | 70.55 |
| 4LOWA | 84 | 149 | -92.75 |
| 4LXLA | 329 | 635 | 344.51 |
| 4LXQA | 273 | 520 | -182.32 |
| 4LZXB | 37 | 60 | 47.87 |
| 4M1GA | 83 | 137 | -202.21 |
| 4M32D | 161 | 312 | -40.48 |
| 4M3LA | 60 | 123 | -220.34 |
| 4M6BC | 23 | 31 | 17.51 |
| 4M70A | 107 | 245 | -155.61 |
| 4MAKA | 73 | 133 | 127.57 |
| 4ME3A | 246 | 438 | -165.99 |
| 4MF5A | 238 | 474 | 432.4 |
| 4MGEB | 107 | 223 | 12.68 |
| 4MHVB | 96 | 179 | -64.81 |
| 4MITF | 35 | 55 | 24.72 |
| 4ML7D | 98 | 153 | 14.07 |
| 4MLIA | 82 | 121 | 49.94 |
| 4MLOA | 267 | 524 | -235.81 |
| 4MMGA | 91 | 173 | -55.47 |
| 4MOYB | 35 | 39 | 15.39 |
| 4MQ3A | 129 | 254 | -51.78 |
| 4MQVB | 26 | 57 | -140.46 |

| | | | |
|---|---|---|---|
| 4MTUA | 143 | 265 | -10.42 |
| 4MUQA | 251 | 472 | 3.33 |
| 4MYYB | 84 | 171 | -141.76 |
| 4MZZB | 32 | 66 | -99.71 |
| 4N1IA | 312 | 547 | 141.63 |
| 4N3BB | 20 | 19 | -0.97 |
| 4N3YA | 35 | 80 | -172.08 |
| 4N67A | 214 | 439 | -85.88 |
| 4N6V5 | 91 | 211 | -57.07 |
| 4N6XA | 91 | 152 | 30.39 |
| 4N7IA | 187 | 329 | 177.92 |
| 4N8MC | 106 | 166 | 1.04 |
| 4N9SA | 295 | 672 | -55.31 |
| 4NAWC | 34 | 68 | -92.12 |
| 4NAZA | 130 | 231 | -147.28 |
| 4NC2A | 121 | 192 | 70.09 |
| 4NCWA | 25 | 34 | 42.45 |
| 4NDIB | 180 | 329 | -125.71 |
| 4NL9C | 60 | 125 | -9.33 |
| 4NM0B | 21 | 32 | -10.66 |
| 4NN2B | 118 | 220 | -166.51 |
| 4NOBA | 112 | 195 | -39.72 |
| 4NPFY | 116 | 282 | -326.44 |
| 4NQWB | 73 | 142 | 112.27 |
| 4NSVA | 264 | 523 | -32.35 |
| 4NUTB | 27 | 55 | -124.31 |
| 4NV4B | 112 | 193 | -41.96 |
| 4NYQA | 153 | 273 | 41.68 |
| 4NZUL | 211 | 354 | -84.91 |
| 4O06A | 102 | 175 | -15.74 |
| 4O1RA | 142 | 251 | 63.93 |
| 4O66A | 67 | 111 | 10.81 |
| 4O6QA | 182 | 350 | 292.24 |
| 4O7JA | 157 | 252 | 121.56 |
| 4OAEA | 160 | 295 | 56.54 |
| 4OCNB | 165 | 296 | 123.44 |
| 4OF6A | 108 | 184 | -106.23 |
| 4OFFA | 124 | 223 | -103.94 |
| 4OGQA | 214 | 488 | 321.87 |
| 4OGQG | 37 | 70 | 80.51 |
| 4OHAA | 242 | 495 | -33.07 |
| 4OIKA | 73 | 118 | -230.53 |
| 4OIYB | 181 | 353 | 82.86 |
| 4OJXA | 363 | 707 | -7.94 |
| 4OKVF | 66 | 140 | -304.39 |
| 4OO4B | 105 | 251 | -125.61 |
| 4OOGA | 109 | 207 | -88.38 |

| | | | |
|---|---|---|---|
| 4OTNA | 123 | 239 | -125.8 |
| 4OUSA | 132 | 223 | -14.97 |
| 4OWFG | 29 | 42 | -87.13 |
| 4OWIA | 33 | 65 | -62.97 |
| 4OWTB | 72 | 116 | 23.93 |
| 4OWWC | 39 | 61 | -53.85 |
| 4OXWA | 106 | 193 | 38.29 |
| 4OXXA | 153 | 306 | 136.73 |
| 4OY3A | 230 | 459 | -26.92 |
| 4OZKA | 49 | 79 | -224.51 |
| 4P0FA | 369 | 680 | -10.34 |
| 4P1ZA | 127 | 246 | -136.4 |
| 4P3AB | 69 | 144 | -396.09 |
| 4P3HA | 190 | 340 | 240.51 |
| 4P3KA | 178 | 330 | 37.26 |
| 4P5AC | 223 | 423 | 66.07 |
| 4PASB | 39 | 86 | -74.27 |
| 4PBZB | 22 | 38 | 32.93 |
| 4PCAB | 218 | 447 | -9.29 |
| 4PDNA | 139 | 260 | 38.57 |
| 4PEOB | 93 | 168 | 22 |
| 4PH1A | 73 | 136 | 0.99 |
| 4PHJA | 173 | 352 | -80.45 |
| 4PHRA | 277 | 559 | -57.05 |
| 4PJ2C | 122 | 259 | -551.02 |
| 4PJEG | 197 | 319 | -125.68 |
| 4PL8H | 26 | 37 | 8.4 |
| 4PMKA | 158 | 290 | -105.79 |
| 4PNDC | 30 | 62 | -73.96 |
| 4PO5C | 162 | 344 | 44.16 |
| 4PP4A | 248 | 483 | -479.08 |
| 4PP8C | 146 | 259 | -202.48 |
| 4PQQA | 156 | 272 | -62.47 |
| 4PS2A | 79 | 135 | 13.17 |
| 4PSFA | 134 | 233 | 69.38 |
| 4PSWC | 38 | 60 | 74.24 |
| 4PUHB | 81 | 149 | 33.02 |
| 4PV2A | 155 | 302 | 242.23 |
| 4PVZC | 35 | 50 | -49.33 |
| 4PWQB | 153 | 265 | -429.7 |
| 4Q0KA | 154 | 274 | 10.79 |
| 4Q4Y4 | 57 | 75 | 92.41 |
| 4Q9BA | 101 | 157 | 9.42 |
| 4QASB | 187 | 353 | 199.82 |
| 4QB0A | 95 | 173 | -46.91 |
| 4QBSA | 138 | 246 | -561.77 |
| 4QF3B | 57 | 87 | -160.2 |

| | | | |
|---|---|---|---|
| 4QICB | 51 | 103 | -57.96 |
| 4QKWB | 99 | 229 | -145.79 |
| 4QLPA | 175 | 345 | 206.82 |
| 4QNCB | 82 | 164 | 44.8 |
| 4QNDA | 97 | 199 | 89.16 |
| 4QPOD | 52 | 103 | -96.26 |
| 4QPTA | 215 | 391 | 31.23 |
| 4QPWA | 142 | 236 | 38.62 |
| 4QQSB | 313 | 549 | 325.49 |
| 4QR9A | 75 | 151 | -183.23 |
| 4QRND | 350 | 736 | 281.53 |
| 4QTQA | 209 | 349 | -16.74 |
| 4QXVA | 116 | 207 | 109.23 |
| 4QXZB | 143 | 257 | -188.14 |
| 4R2QA | 88 | 160 | -321.3 |
| 4R2YD | 62 | 101 | -261.39 |
| 4R38A | 109 | 201 | -58.08 |
| 4R3HA | 157 | 290 | -32.44 |
| 4R3QA | 69 | 144 | -249.34 |
| 4R5RB | 68 | 120 | -468.71 |
| 4R6FA | 329 | 686 | -268.13 |
| 4R6RG | 133 | 225 | 238.78 |
| 4R7AB | 366 | 643 | -63.96 |
| 4R8TA | 21 | 24 | 3.47 |
| 4R9KB | 155 | 301 | 205.28 |
| 4RA6A | 177 | 344 | 221.48 |
| 4RAXA | 227 | 405 | 141.8 |
| 4RAYB | 133 | 239 | -9.24 |
| 4RDJA | 293 | 497 | 578.63 |
| 4RE1B | 202 | 358 | -215.44 |
| 4RE1C | 50 | 80 | 70.92 |
| 4REHA | 149 | 272 | -25.62 |
| 4REKA | 499 | 964 | 506.26 |
| 4REYB | 22 | 27 | 28.99 |
| 4RFUB | 117 | 195 | 139.39 |
| 4RGDB | 70 | 148 | 127.34 |
| 4RIQU | 23 | 47 | 3.1 |
| 4RJFB | 21 | 25 | -17.99 |
| 4RKHF | 46 | 73 | -248.96 |
| 4RLCA | 135 | 230 | 54.21 |
| 4RLRA | 126 | 258 | -121.4 |
| 4ROWA | 191 | 358 | -283.08 |
| 4RQRA | 107 | 195 | -99.2 |
| 4RRFA | 138 | 257 | -93.41 |
| 4RT4E | 21 | 50 | -36.85 |
| 4RU3A | 205 | 345 | 174.52 |
| 4RUVA | 106 | 198 | 36.58 |

| | | | |
|---|---|---|---|
| 4RVQA | 122 | 255 | -242.09 |
| 4RWWB | 106 | 165 | -54.86 |
| 4RYOA | 151 | 314 | 79.53 |
| 4S1BA | 213 | 442 | -34.93 |
| 4SGBI | 51 | 87 | -243.13 |
| 4TKBA | 133 | 242 | -108.26 |
| 4TPSC | 140 | 263 | -153.4 |
| 4TQ1B | 32 | 57 | -125.14 |
| 4TSDA | 179 | 328 | -125.66 |
| 4TTLA | 22 | 39 | -166.35 |
| 4TTWA | 132 | 220 | -76.74 |
| 4TXRA | 159 | 347 | -135.67 |
| 4U0YA | 75 | 145 | 61.08 |
| 4U5HC | 84 | 169 | -616.75 |
| 4U6OB | 156 | 267 | 88.3 |
| 4U7IA | 93 | 199 | -70.4 |
| 4U88A | 93 | 171 | -181.53 |
| 4U9HL | 533 | 1087 | 316.61 |
| 4U9HS | 265 | 500 | 383.45 |
| 4U9WD | 192 | 369 | -120.45 |
| 4UE0A | 115 | 176 | 50.64 |
| 4UE8B | 37 | 60 | 86.52 |
| 4UE9B | 36 | 51 | -31.25 |
| 4UHDA | 274 | 574 | 333.18 |
| 4UHQB | 134 | 270 | 229.58 |
| 4UHTB | 102 | 183 | 30.67 |
| 4UI1D | 72 | 132 | -220.43 |
| 4UJ0B | 47 | 86 | -333.52 |
| 4UN2B | 43 | 83 | -13.88 |
| 4UP0A | 85 | 151 | -144.11 |
| 4UQWA | 163 | 337 | 105.63 |
| 4UQZB | 24 | 25 | 32.07 |
| 4UR6B | 62 | 109 | 260.25 |
| 4USGB | 66 | 115 | 71.39 |
| 4UU3B | 164 | 296 | -2.49 |
| 4UWWA | 130 | 242 | -270.84 |
| 4UXUA | 210 | 336 | -127.83 |
| 4UYPA | 144 | 254 | 255.77 |
| 4UYRA | 188 | 331 | -455.25 |
| 4UZ3C | 47 | 77 | 60.47 |
| 4UZZB | 65 | 131 | -89.5 |
| 4V0WD | 22 | 24 | 8.89 |
| 4V3IA | 153 | 320 | -85.16 |
| 4V3KF | 70 | 121 | -251.58 |
| 4W8HA | 127 | 238 | 39.37 |
| 4W8PB | 21 | 44 | -13.57 |
| 4WCGA | 61 | 118 | 74.23 |

| | | | |
|---|---|---|---|
| 4WEEA | 135 | 227 | 179.62 |
| 4WESB | 458 | 934 | 461.21 |
| 4WFCC | 93 | 194 | -174.96 |
| 4WFTA | 94 | 174 | -66.05 |
| 4WH9A | 177 | 334 | 110.03 |
| 4WHIA | 102 | 163 | 177.01 |
| 4WHSF | 238 | 412 | 292.36 |
| 4WI1A | 440 | 861 | -464.03 |
| 4WNDB | 26 | 32 | 4.36 |
| 4WOHA | 158 | 318 | -18.26 |
| 4WOLC | 30 | 59 | 74.71 |
| 4WSPA | 79 | 139 | 11.29 |
| 4WULB | 64 | 133 | -108.03 |
| 4WV4B | 93 | 186 | 23.1 |
| 4WWBA | 112 | 246 | -60.08 |
| 4WWMA | 79 | 143 | -45.72 |
| 4WX4A | 201 | 393 | -140.56 |
| 4WXAC | 78 | 149 | 55.64 |
| 4WXAE | 69 | 137 | -11.31 |
| 4WY4D | 65 | 139 | -58.39 |
| 4WZXA | 87 | 185 | -161.68 |
| 4X08A | 133 | 248 | -197.28 |
| 4X1JA | 83 | 126 | -352.67 |
| 4X2OA | 179 | 313 | 54.32 |
| 4X33A | 60 | 96 | -140.04 |
| 4X3SA | 61 | 107 | 59.77 |
| 4X5LA | 96 | 153 | 137.04 |
| 4X8KB | 32 | 64 | -95.61 |
| 4X9CA | 57 | 97 | 11.36 |
| 4X9ZB | 46 | 65 | -366.96 |
| 4XBAA | 199 | 398 | -106.2 |
| 4XDIB | 123 | 255 | 19.66 |
| 4XDQA | 237 | 414 | 313.75 |
| 4XEZA | 186 | 348 | -268.26 |
| 4XHTA | 94 | 181 | -66.94 |
| 4XINB | 114 | 185 | 54.35 |
| 4XIZN | 70 | 137 | -137.43 |
| 4XKLB | 30 | 50 | -80.79 |
| 4XKZA | 251 | 500 | -152.97 |
| 4XLGB | 67 | 142 | -129.73 |
| 4XMRB | 252 | 480 | -31.61 |
| 4XRMB | 64 | 122 | -27.3 |
| 4XTRG | 22 | 42 | -68.2 |
| 4XW3B | 194 | 329 | -85.15 |
| 4XZFA | 121 | 214 | 21.43 |
| 4Y0GA | 82 | 133 | 77.68 |
| 4Y6WA | 211 | 454 | -365.03 |

| | | | |
|---|---|---|---|
| 4Y88A | 108 | 188 | 62.72 |
| 4Y9VA | 603 | 1179 | -37.77 |
| 4Y9WA | 328 | 602 | 238.3 |
| 4YDXA | 67 | 121 | -46.95 |
| 4YEPB | 185 | 316 | -48.38 |
| 4YG0A | 164 | 283 | 97.03 |
| 4YGBD | 63 | 130 | -29.73 |
| 4YH8B | 55 | 90 | 19.18 |
| 4YI0A | 27 | 30 | -56.62 |
| 4YIZF | 39 | 65 | 40.58 |
| 4YK2A | 107 | 221 | -105.41 |
| 4YKIB | 255 | 477 | -166.28 |
| 4YMIA | 178 | 327 | 173.27 |
| 4YMYA | 153 | 261 | 157.67 |
| 4YNHA | 58 | 118 | 46.07 |
| 4YSXG | 155 | 311 | 220.81 |
| 4YTKA | 116 | 198 | 23.55 |
| 4YV4B | 51 | 99 | -139.49 |
| 4YWCD | 22 | 46 | -17.85 |
| 4YZ0A | 198 | 405 | -121.58 |
| 4Z02B | 104 | 164 | 12.61 |
| 4Z04A | 124 | 212 | 159.15 |
| 4Z3CC | 45 | 70 | -189.59 |
| 4Z3HA | 197 | 316 | 140.8 |
| 4Z47A | 196 | 387 | 17.61 |
| 4Z4AB | 75 | 125 | -206.56 |
| 4Z80D | 31 | 52 | -4.44 |
| 4Z9VE | 21 | 28 | -19.82 |
| 4ZAVA | 208 | 400 | 111.94 |
| 4ZBHA | 119 | 189 | 163.8 |
| 4ZBJD | 21 | 28 | 15.46 |
| 4ZC3A | 57 | 111 | -4.71 |
| 4ZDTA | 69 | 117 | -204.31 |
| 4ZDTB | 69 | 140 | -70.63 |
| 4ZEQB | 26 | 50 | 22.88 |
| 4ZFOK | 35 | 58 | -232.27 |
| 4ZGMA | 100 | 179 | -218.62 |
| 4ZHNA | 201 | 356 | 93.05 |
| 4ZORE | 129 | 216 | 166.94 |
| 4ZOSD | 98 | 176 | -38.62 |
| 4ZQKB | 106 | 169 | -21.21 |
| 4ZQXA | 237 | 403 | 174.09 |
| 4ZRKF | 21 | 40 | -49.04 |
| 5A0LA | 205 | 394 | 92.78 |
| 5A0NA | 214 | 417 | -86.4 |
| 5A27A | 411 | 760 | 276.06 |
| 5A53B | 22 | 22 | 43.24 |

| | | | |
|---|---|---|---|
| 5A6WC | 83 | 120 | -18.57 |
| 5ABVH | 59 | 96 | 24.89 |
| 5ABYF | 39 | 66 | 72.69 |
| 5AEAA | 100 | 170 | -9.52 |
| 5AEJB | 113 | 175 | -225.23 |
| 5AF0A | 237 | 470 | -162.68 |
| 5AFDA | 300 | 609 | 107.07 |
| 5AIGB | 124 | 249 | 49 |
| 5AJJA | 138 | 302 | -62.51 |
| 5AKRA | 337 | 614 | 619.16 |
| 5AL6A | 44 | 98 | -201.57 |
| 5AMHA | 106 | 180 | -36.64 |
| 5AN5B | 21 | 41 | 0.49 |
| 5AN5E | 24 | 43 | -29.95 |
| 5ANRC | 23 | 29 | 43.09 |
| 5AOND | 49 | 105 | -2.18 |
| 5AOTA | 102 | 169 | -61.2 |
| 5AQ0A | 81 | 133 | 145.08 |
| 5AQCA | 183 | 373 | -29.35 |
| 5ARMA | 127 | 249 | -845.29 |
| 5AYSC | 109 | 206 | -107.2 |
| 5B0GA | 100 | 177 | -40.46 |
| 5B0HB | 133 | 236 | 30.76 |
| 5B1AU | 79 | 146 | -166.52 |
| 5B1AW | 58 | 120 | -20.88 |
| 5B1AX | 49 | 83 | 89.38 |
| 5B1AZ | 43 | 79 | 92.51 |
| 5B1NA | 59 | 130 | 14.94 |
| 5B1RA | 116 | 200 | -225.95 |
| 5B5HA | 223 | 440 | 471.95 |
| 5B5IB | 68 | 143 | -297.37 |
| 5B66F | 33 | 72 | 72.47 |
| 5B71E | 93 | 133 | 116.46 |
| 5B82A | 161 | 308 | -47.81 |
| 5B8DA | 100 | 172 | -86.93 |
| 5B8FB | 158 | 307 | 192.88 |
| 5BOWA | 151 | 250 | 35.86 |
| 5BR4A | 385 | 811 | 503.18 |
| 5BRKB | 64 | 141 | -198.84 |
| 5BW0D | 84 | 140 | -62.85 |
| 5BWDC | 39 | 55 | 9.59 |
| 5BXVD | 33 | 58 | 97.48 |
| 5BY1A | 71 | 156 | -13.29 |
| 5BY8B | 85 | 131 | 219.23 |
| 5C0ZC | 104 | 190 | -21.62 |
| 5C10A | 230 | 446 | -13.39 |
| 5C17A | 206 | 405 | -5.77 |

| | | | |
|---|---|---|---|
| 5C5TB | 179 | 331 | -217.95 |
| 5C6HT | 26 | 49 | 14.45 |
| 5CBAF | 64 | 96 | -106.49 |
| 5CEGC | 85 | 162 | -11.96 |
| 5CESB | 91 | 163 | -8.48 |
| 5CGQA | 268 | 543 | 402.27 |
| 5CGQB | 390 | 803 | 349.86 |
| 5CJ9A | 146 | 289 | -14.8 |
| 5CKLA | 181 | 376 | -149.43 |
| 5COYB | 65 | 109 | -137.92 |
| 5CQ2A | 76 | 119 | 102.2 |
| 5CQEA | 158 | 316 | 73.88 |
| 5CR8D | 113 | 179 | -69.35 |
| 5CTVA | 176 | 352 | 157.96 |
| 5CWTC | 40 | 87 | -214.96 |
| 5CWWC | 32 | 58 | -51.73 |
| 5CX3E | 20 | 27 | 36.89 |
| 5CX3F | 25 | 37 | -33.28 |
| 5CX7P | 138 | 254 | 102.4 |
| 5CYAA | 161 | 311 | -350.04 |
| 5CYVA | 161 | 311 | -350.04 |
| 5D14A | 70 | 125 | -214.24 |
| 5D18A | 202 | 435 | -38.44 |
| 5D1LB | 111 | 162 | -210.6 |
| 5D22B | 121 | 223 | -175.77 |
| 5D2MG | 21 | 21 | 59.24 |
| 5D3XB | 141 | 262 | -378.35 |
| 5D50L | 104 | 188 | 120.82 |
| 5D60B | 61 | 145 | -235.02 |
| 5D8VA | 83 | 169 | 41.26 |
| 5DCUA | 364 | 703 | -107.32 |
| 5DHVN | 58 | 113 | -102.59 |
| 5DLOA | 115 | 200 | -25.69 |
| 5DOCA | 197 | 371 | -205.06 |
| 5DOMA | 90 | 197 | -356.88 |
| 5DOWH | 20 | 41 | -30.34 |
| 5DP2A | 335 | 642 | 373.07 |
| 5DQSD | 87 | 161 | 156.02 |
| 5DRKC | 89 | 145 | 129.04 |
| 5DUSA | 165 | 330 | 132.12 |
| 5DVIA | 396 | 797 | 294.15 |
| 5E0YA | 68 | 121 | 136.76 |
| 5E1DB | 225 | 438 | -43.22 |
| 5E1YA | 95 | 162 | 21.83 |
| 5E9EB | 261 | 495 | 20.56 |
| 5ECWB | 123 | 247 | 73 |
| 5EH6A | 28 | 54 | 81.89 |

| | | | |
|---|---|---|---|
| 5EJDO | 74 | 139 | 91.77 |
| 5EL9A | 199 | 379 | -56.2 |
| 5EMXA | 53 | 104 | -80.96 |
| 5EN2C | 141 | 250 | -117.22 |
| 5EO6B | 307 | 591 | -14.3 |
| 5EOAA | 72 | 165 | -281.21 |
| 5EP6A | 36 | 77 | -140.71 |
| 5EPMD | 33 | 58 | -261.54 |
| 5ET0D | 20 | 28 | -2.37 |
| 5ET1D | 23 | 37 | -16.15 |
| 5ET3B | 30 | 61 | -109.46 |
| 5EVCA | 237 | 477 | 214.39 |
| 5EWOA | 214 | 342 | 139.21 |
| 5F3BD | 103 | 159 | -192.75 |
| 5F4CB | 91 | 176 | 42.17 |
| 5F4QA | 197 | 373 | -680.41 |
| 5F5OF | 27 | 45 | 45.67 |
| 5F74B | 20 | 35 | -44.69 |
| 5FAFA | 113 | 203 | -5.69 |
| 5FCGC | 26 | 53 | -48.36 |
| 5FEBA | 122 | 227 | -122.26 |
| 5FEYB | 79 | 131 | -244.71 |
| 5FFDA | 137 | 282 | -200.49 |
| 5FFXA | 138 | 298 | -242.68 |
| 5FI3B | 344 | 644 | 408.91 |
| 5FJDD | 111 | 237 | -479.16 |
| 5FMKB | 21 | 40 | 27.6 |
| 5FMLB | 196 | 388 | -1.62 |
| 5FMNA | 86 | 182 | -132.5 |
| 5FO8C | 129 | 188 | -54.32 |
| 5FS4A | 118 | 192 | 14.57 |
| 5FVDD | 27 | 47 | -28.4 |
| 5FVKC | 21 | 26 | 91.89 |
| 5FVLC | 24 | 41 | 94.31 |
| 5FW5C | 23 | 34 | 29.53 |
| 5FYDA | 259 | 514 | 178.83 |
| 5FZTB | 23 | 43 | 8.36 |
| 5G0RC | 248 | 462 | 103.49 |
| 5G3YA | 213 | 427 | 49.26 |
| 5G5HA | 173 | 305 | -43.73 |
| 5GGBA | 290 | 543 | 134.52 |
| 5GGNB | 151 | 283 | 54.4 |
| 5GHWP | 21 | 41 | -64.67 |
| 5GJIA | 107 | 203 | -61.91 |
| 5GJKB | 68 | 115 | 64.75 |
| 5GK9B | 24 | 28 | 27.47 |
| 5GM9A | 213 | 409 | -339.83 |

| | | | |
|---|---|---|---|
| 5GNAB | 56 | 117 | -50.1 |
| 5GPIC | 37 | 44 | 3.39 |
| 5GQIA | 246 | 459 | 167.31 |
| 5GRMB | 190 | 360 | -53.75 |
| 5GRQB | 85 | 180 | -146.73 |
| 5GRQC | 30 | 54 | -12.34 |
| 5GTUB | 24 | 49 | -30.24 |
| 5GU8A | 71 | 108 | 87.6 |
| 5GXWB | 27 | 33 | 43.56 |
| 5GY7A | 338 | 670 | 306.21 |
| 5GYQA | 169 | 283 | 74.38 |
| 5H17B | 27 | 48 | -70.21 |
| 5H3JB | 21 | 25 | -91.28 |
| 5H3VB | 103 | 166 | 97.12 |
| 5H4GB | 139 | 290 | 112.44 |
| 5H7YB | 25 | 52 | -31.4 |
| 5H8JP | 279 | 504 | -21.61 |
| 5H94B | 97 | 149 | -66.92 |
| 5HA6A | 93 | 198 | -253.26 |
| 5HB6B | 143 | 251 | 263.01 |
| 5HB8A | 109 | 184 | 142.23 |
| 5HDKC | 96 | 182 | 1.31 |
| 5HDMA | 428 | 851 | 711.57 |
| 5HFSB | 65 | 98 | 127.92 |
| 5HHJA | 294 | 572 | -50.99 |
| 5HI4A | 89 | 131 | -110.83 |
| 5HI8B | 136 | 236 | -131.96 |
| 5HQWA | 287 | 535 | 192.64 |
| 5HSFB | 108 | 178 | -149.98 |
| 5HT2A | 255 | 482 | -168.28 |
| 5HT6A | 123 | 198 | 42.06 |
| 5HT8A | 87 | 159 | 49.33 |
| 5HVZC | 23 | 42 | -60.9 |
| 5HWAA | 259 | 542 | -73.33 |
| 5HXLA | 123 | 209 | -65.8 |
| 5HYCC | 29 | 29 | 64.85 |
| 5I0QA | 34 | 67 | -121.8 |
| 5I29A | 139 | 294 | -90.99 |
| 5I2QA | 105 | 206 | -6.59 |
| 5I32A | 238 | 511 | 581.56 |
| 5I4ZA | 74 | 149 | -199.69 |
| 5I5NB | 136 | 269 | -151.19 |
| 5I8TA | 179 | 315 | -102.6 |
| 5IBNA | 111 | 230 | 32.68 |
| 5IBWC | 43 | 91 | -86.99 |
| 5ICVA | 180 | 344 | -82.54 |
| 5II0A | 99 | 170 | -175.7 |

| | | | |
|---|---|---|---|
| 5IJHA | 181 | 355 | -255.8 |
| 5IL3A | 536 | 1098 | -110.77 |
| 5IMMA | 120 | 202 | -140.18 |
| 5IO8B | 101 | 206 | -87.92 |
| 5IO9B | 95 | 164 | -17.22 |
| 5IQNA | 132 | 222 | 36.81 |
| 5IXHB | 161 | 284 | 230.19 |
| 5J1SB | 235 | 451 | -142.23 |
| 5J4FB | 97 | 154 | -54.88 |
| 5J4SB | 67 | 112 | -520.54 |
| 5J6YA | 188 | 341 | 99.31 |
| 5J7DE | 106 | 200 | -65.68 |
| 5JA9C | 101 | 188 | -91.15 |
| 5JBSB | 126 | 225 | 201.4 |
| 5JBTY | 38 | 61 | -114.51 |
| 5JDAA | 359 | 701 | -141.35 |
| 5JFFB | 51 | 98 | -173.95 |
| 5JG9B | 47 | 84 | -435.42 |
| 5JGEE | 30 | 56 | -105.56 |
| 5JLBA | 248 | 429 | -278.7 |
| 5JNBG | 28 | 46 | 20.38 |
| 5JNVA | 154 | 323 | -235.79 |
| 5JOEA | 92 | 153 | -150.85 |
| 5JQAB | 20 | 40 | -5.63 |
| 5JQFB | 21 | 31 | 85.84 |
| 5JQIB | 277 | 450 | 318.17 |
| 5JSVA | 370 | 735 | 207.46 |
| 5JUGA | 168 | 343 | 47.08 |
| 5JZED | 76 | 142 | -123.44 |
| 5K2ME | 52 | 73 | 44.76 |
| 5K3QC | 42 | 71 | -283.33 |
| 5K6DA | 77 | 142 | -365.5 |
| 5K8QB | 26 | 50 | 11.44 |
| 5K9BA | 179 | 364 | 240.51 |
| 5KC1C | 33 | 61 | -13.25 |
| 5KLCA | 93 | 159 | 46.85 |
| 5KN8A | 222 | 452 | 111.69 |
| 5KO4B | 101 | 204 | 32.4 |
| 5KO6A | 285 | 543 | 200 |
| 5KOED | 454 | 847 | -235.43 |
| 5KP7B | 79 | 147 | 78.07 |
| 5KSPB | 155 | 276 | -100.73 |
| 5KWYD | 132 | 216 | -104.07 |
| 5KXHB | 38 | 63 | -270.4 |
| 5KYEC | 76 | 141 | 28.53 |
| 5L04B | 37 | 42 | 93.86 |
| 5L16A | 323 | 595 | 239.34 |

| | | | |
|---|---|---|---|
| 5L2LA | 72 | 109 | -208.46 |
| 5L37C | 86 | 134 | 197.28 |
| 5L71A | 165 | 319 | -68.28 |
| 5L8LB | 109 | 184 | -140.29 |
| 5LCYC | 89 | 198 | -86.15 |
| 5LI1B | 20 | 27 | -4.83 |
| 5LJMA | 203 | 427 | -61.6 |
| 5LKVA | 137 | 290 | -104.55 |
| 5LLJB | 57 | 83 | 2.91 |
| 5LP0A | 140 | 314 | -264.32 |
| 5LPIC | 134 | 245 | -255.07 |
| 5LQFB | 264 | 556 | 121.52 |
| 5LRRF | 226 | 426 | -259.67 |
| 5LS7B | 128 | 225 | -99.11 |
| 5LSLE | 62 | 103 | 186.92 |
| 5LSWD | 21 | 39 | -4.84 |
| 5LTEA | 333 | 642 | 196.5 |
| 5LUSI | 50 | 102 | -87.25 |
| 5LXRB | 40 | 66 | 182.92 |
| 5LXXA | 477 | 998 | 263.98 |
| 5LY8A | 230 | 406 | 243.46 |
| 5M33B | 86 | 165 | -366.82 |
| 5M5TB | 360 | 618 | 146.47 |
| 5M72B | 21 | 32 | 15.29 |
| 5ME5A | 152 | 283 | -133.12 |
| 5MFOC | 199 | 432 | -292.3 |
| 5MJRA | 186 | 392 | 13.06 |
| 5MK0B | 20 | 32 | -34.62 |
| 5MKWB | 117 | 229 | -157.93 |
| 5ML3B | 149 | 262 | -54.15 |
| 5MR1A | 101 | 164 | -58.92 |
| 5MSMB | 132 | 198 | 79.93 |
| 5MSMC | 27 | 31 | -5.43 |
| 5MU3C | 24 | 39 | -21.57 |
| 5MU9A | 292 | 538 | 310.7 |
| 5MVHA | 397 | 697 | 18.85 |
| 5MVWC | 37 | 83 | -145.76 |
| 5N4CF | 32 | 45 | 29.19 |
| 5N4KA | 130 | 200 | 186.94 |
| 5N6YF | 111 | 222 | -189.54 |
| 5NAJC | 111 | 196 | -72.54 |
| 5NCWB | 33 | 43 | 30.04 |
| 5NDCC | 31 | 60 | 9.3 |
| 5NGNA | 32 | 62 | -537.77 |
| 5NHUJ | 29 | 44 | 132.1 |
| 5NIHA | 262 | 490 | 55.77 |
| 5NQFB | 35 | 47 | 7.69 |

| | | | |
|---|---|---|---|
| 5NVLD | 67 | 106 | 29.2 |
| 5NWPB | 197 | 335 | 534.74 |
| 5O1XA | 162 | 298 | 98.6 |
| 5O45A | 129 | 221 | -227.2 |
| 5O5SA | 123 | 250 | 79.76 |
| 5O63A | 159 | 281 | 162.82 |
| 5O6TB | 63 | 102 | -218.49 |
| 5O99B | 59 | 93 | 166 |
| 5OD4A | 124 | 188 | 151.18 |
| 5ODKA | 173 | 313 | 42.17 |
| 5OHOB | 113 | 192 | -58 |
| 5OWUB | 29 | 42 | 79.26 |
| 5PAGA | 57 | 100 | -146.26 |
| 5PXLB | 177 | 344 | -565.3 |
| 5SUYC | 95 | 154 | 3.97 |
| 5SVXA | 49 | 77 | -3.49 |
| 5SXPG | 21 | 21 | 168.59 |
| 5SZDB | 70 | 119 | -23.56 |
| 5T0FB | 32 | 59 | 20.73 |
| 5T3UA | 136 | 258 | -7.36 |
| 5T46D | 36 | 57 | 14.11 |
| 5T48B | 36 | 65 | -58.89 |
| 5T51B | 90 | 193 | -348.01 |
| 5T59C | 32 | 51 | 10.67 |
| 5T7AA | 93 | 149 | 28.12 |
| 5TDBA | 70 | 117 | -102.31 |
| 5TDRA | 70 | 123 | -185.33 |
| 5TDYC | 42 | 85 | -38.66 |
| 5TEYB | 241 | 426 | -169.06 |
| 5TGIC | 20 | 26 | -3.02 |
| 5TGID | 20 | 29 | -7.58 |
| 5TIFA | 182 | 389 | 33.01 |
| 5TP1A | 150 | 274 | -55.78 |
| 5TP1S | 20 | 27 | 3.04 |
| 5TPKA | 202 | 333 | 203.33 |
| 5TRBA | 69 | 127 | -254.4 |
| 5TUUA | 144 | 278 | -154.97 |
| 5TVIV | 90 | 179 | -223.09 |
| 5TWAD | 25 | 49 | 16.24 |
| 5TWBA | 344 | 653 | -34.39 |
| 5TX4B | 92 | 142 | -214.56 |
| 5TZGA | 75 | 154 | 43.36 |
| 5TZQA | 132 | 271 | -49.89 |
| 5TZQC | 22 | 46 | -77.83 |
| 5U23B | 363 | 746 | -556.48 |
| 5U9NA | 125 | 207 | -237.24 |
| 5UD5A | 86 | 155 | -49.22 |

| | | | |
|---|---|---|---|
| 5UEJA | 375 | 737 | 173.94 |
| 5UFYA | 179 | 364 | 1.71 |
| 5UJCA | 267 | 502 | 215.91 |
| 5UN7B | 61 | 96 | -31.34 |
| 5URVB | 129 | 239 | -275.37 |
| 5UUKA | 152 | 303 | 6.95 |
| 5UULB | 23 | 45 | -55.12 |
| 5V1AB | 26 | 35 | 36.07 |
| 5V1YA | 114 | 195 | -148.36 |
| 5V2CH | 65 | 122 | 68.74 |
| 5V2CJ | 39 | 75 | 137.28 |
| 5V2CL | 37 | 65 | -35.38 |
| 5V50C | 129 | 235 | -254.81 |
| 5V77A | 106 | 173 | 68.79 |
| 5V87A | 147 | 299 | -267.21 |
| 5V89C | 67 | 132 | -137.06 |
| 5V8KB | 25 | 51 | 52.88 |
| 5VCYA | 296 | 548 | 210.96 |
| 5VHGA | 150 | 300 | 14.83 |
| 5VLAZ | 24 | 40 | 19.4 |
| 5VMDD | 66 | 106 | -274.79 |
| 5VMRC | 38 | 75 | -102.82 |
| 5VT9C | 30 | 50 | 46.96 |
| 5VZ3A | 108 | 175 | -243.08 |
| 5W9QA | 56 | 87 | -294.68 |
| 5WFBA | 132 | 283 | -21.6 |
| 5WHTA | 115 | 207 | -42.32 |
| 5WHXB | 253 | 511 | -211.06 |
| 5WO2B | 89 | 200 | -248.81 |
| 5WS7B | 156 | 304 | -15.41 |
| 5WSVB | 29 | 62 | 27.14 |
| 5WTQC | 105 | 163 | -40.6 |
| 5WWOC | 46 | 82 | -28.89 |
| 5WWWA | 94 | 180 | -109.45 |
| 5WXMU | 27 | 55 | 14.7 |
| 5X5JA | 118 | 224 | -13.32 |
| 5X8IA | 328 | 649 | -117.11 |
| 5XA5B | 27 | 47 | 12.44 |
| 5XADC | 24 | 32 | 25.41 |
| 5XAUC | 70 | 136 | -126.63 |
| 5XCTA | 167 | 316 | -56.7 |
| 5XECA | 82 | 144 | 99.21 |
| 5XG5A | 145 | 245 | 168.19 |
| 5XJGB | 30 | 33 | 13.66 |
| 5XTRC | 245 | 510 | -420.29 |
| 5YGEA | 169 | 305 | 75.99 |
| 6ANWA | 96 | 172 | 92.67 |

| | | | |
|---|---|---|---|
| 6AO7A | 153 | 276 | -63.92 |
| 6B0GE | 154 | 255 | -806.77 |
| 6B0SC | 65 | 113 | -120.62 |
| 6B29A | 57 | 94 | 58.4 |
| 6B4AA | 92 | 161 | 52.73 |
| 8A3HA | 300 | 623 | -70.21 |
