## Supplementary-Data-D for "Estimating Boltzmann Statistical Energy for Proteins from 3D Structure Using Pairwise Amino Acid Energy Matrix - GEM-X2120-N871"

| PDB | No of aa's | No of ints. | X2120-N871 dataset |
|---|---|---|---|
| 1A11A | 25 | 60 | -51.21 |
| 1AA3A | 63 | 157 | -52.76 |
| 1AIWA | 62 | 120 | -59.43 |
| 1ANSA | 27 | 55 | -197.58 |
| 1APFA | 49 | 99 | -149.24 |
| 1AUUB | 55 | 108 | 3.39 |
| 1AXHA | 37 | 72 | -288.1 |
| 1B22A | 70 | 162 | 89.46 |
| 1B4GA | 29 | 45 | -5.79 |
| 1BCTA | 69 | 147 | 141.89 |
| 1BDEA | 33 | 75 | -51.16 |
| 1BH7A | 31 | 46 | 48.83 |
| 1BHIA | 38 | 66 | -71.43 |
| 1BL1A | 31 | 43 | -15.74 |
| 1BM4A | 32 | 51 | 61.65 |
| 1BNXA | 33 | 60 | 116.19 |
| 1BOMA | 20 | 37 | -117.71 |
| 1BONB | 27 | 48 | -31.5 |
| 1BQFA | 25 | 36 | 0.45 |
| 1BTRA | 20 | 38 | 22.33 |
| 1BTTA | 21 | 43 | 37.08 |
| 1BW6A | 56 | 107 | 34.44 |
| 1BYYA | 21 | 36 | 0.08 |
| 1C01A | 76 | 151 | -173.38 |
| 1C6WA | 33 | 68 | -338.71 |
| 1CFFB | 20 | 38 | -25.64 |
| 1CFHA | 47 | 82 | -140.67 |
| 1CKYA | 26 | 33 | 30.69 |
| 1CLDA | 33 | 55 | -248.11 |
| 1CO4A | 42 | 75 | -71 |
| 1CQ0A | 28 | 61 | 11.32 |
| 1CZ6A | 25 | 40 | -148.53 |
| 1D1HA | 35 | 68 | -474.82 |
| 1D9OA | 20 | 31 | 15.38 |
| 1DENA | 60 | 102 | -246.55 |
| 1DKCA | 38 | 67 | -211.97 |
| 1DSQA | 21 | 36 | -69.54 |
| 1DSVA | 31 | 58 | 4.64 |
| 1DWLA | 59 | 116 | -62.43 |
| 1DXZA | 32 | 74 | -14.48 |
| 1E0LA | 37 | 65 | 18.59 |
| 1E0NA | 27 | 37 | 21.54 |
| 1E4TA | 37 | 58 | -258.4 |
| 1E8QA | 46 | 90 | -191.01 |
| 1ECIA | 37 | 71 | -36.95 |
| 1ED7A | 45 | 91 | 10.34 |

| | | | |
|---|---|---|---|
| 1EDWA | 26 | 52 | 89.69 |
| 1EITA | 36 | 59 | -443.18 |
| 1EJQB | 28 | 29 | 50.28 |
| 1EMZA | 21 | 42 | 45.16 |
| 1ETGB | 23 | 54 | -129.91 |
| 1EV0B | 58 | 108 | 40.61 |
| 1EWSA | 32 | 62 | -317.89 |
| 1EZEA | 38 | 82 | -3.7 |
| 1FACA | 21 | 31 | -11.55 |
| 1FAFA | 79 | 174 | -86.64 |
| 1FARA | 52 | 83 | -214.63 |
| 1FCTA | 32 | 68 | 119.42 |
| 1FDFA | 25 | 51 | 38.75 |
| 1FEXA | 59 | 115 | -69.8 |
| 1FGEA | 20 | 31 | 8.21 |
| 1FI0A | 21 | 53 | -26.16 |
| 1FMHB | 31 | 71 | -45.71 |
| 1FREA | 39 | 83 | -273.73 |
| 1FRYA | 29 | 46 | 71.89 |
| 1FU9A | 36 | 67 | -112.47 |
| 1FVYA | 31 | 60 | -63.36 |
| 1FW5A | 20 | 50 | 11.84 |
| 1FWOA | 35 | 55 | -94.58 |
| 1G1ZA | 31 | 48 | -213.01 |
| 1GEAA | 20 | 48 | 11.13 |
| 1GURA | 34 | 55 | -137.46 |
| 1GYAA | 105 | 201 | -179.51 |
| 1H8BB | 23 | 51 | -15.47 |
| 1HCCA | 59 | 90 | 8.11 |
| 1HCEA | 118 | 207 | -434.56 |
| 1HD6A | 37 | 85 | -232.63 |
| 1HDJA | 77 | 154 | -8.93 |
| 1HJIB | 26 | 56 | -64.22 |
| 1HN3A | 40 | 62 | 23.86 |
| 1HO7A | 20 | 45 | 22.62 |
| 1HP3A | 32 | 52 | -269.88 |
| 1HP9A | 22 | 47 | -381.03 |
| 1HZNA | 29 | 42 | 22.56 |
| 1I25A | 37 | 50 | -215.4 |
| 1IIEC | 75 | 169 | -63.09 |
| 1IIJA | 35 | 63 | -6.2 |
| 1IMTA | 80 | 147 | -305.95 |
| 1IYCA | 36 | 66 | 36.51 |
| 1J5LA | 30 | 49 | -384.82 |
| 1JBIA | 100 | 183 | 8.17 |
| 1JEIA | 53 | 104 | 8.11 |
| 1JGNB | 22 | 27 | 42.41 |

| | | | |
|---|---|---|---|
| 1JH4B | 22 | 29 | 77.49 |
| 1JO6A | 45 | 69 | -59.81 |
| 1JSPA | 19 | 22 | 5.78 |
| 1JUNB | 43 | 113 | -159.27 |
| 1JY9A | 19 | 27 | 3.28 |
| 1JZPA | 20 | 40 | -68.08 |
| 1K18A | 31 | 59 | -22.62 |
| 1K4UP | 32 | 67 | 144.63 |
| 1KALA | 29 | 54 | -162.68 |
| 1KBFA | 49 | 82 | -207.69 |
| 1KCNA | 21 | 35 | -35.41 |
| 1KDLA | 23 | 47 | -22.22 |
| 1KFTA | 56 | 155 | 73.28 |
| 1KN6A | 73 | 158 | -49.67 |
| 1KQKA | 80 | 146 | -103.24 |
| 1KV4A | 42 | 90 | 146.72 |
| 1LBJA | 22 | 49 | -25.65 |
| 1LFCA | 25 | 46 | -112.01 |
| 1LIQA | 27 | 54 | -116.28 |
| 1LJZB | 25 | 40 | -36.66 |
| 1LKQA | 21 | 34 | -123.1 |
| 1LMRA | 35 | 55 | -198.95 |
| 1LOIA | 25 | 51 | 16.4 |
| 1LV4A | 21 | 29 | 64.57 |
| 1LV9A | 63 | 118 | -37.34 |
| 1LYPA | 32 | 67 | -48.32 |
| 1M4EA | 20 | 31 | -349.5 |
| 1M9OA | 40 | 78 | -49.36 |
| 1MEAA | 28 | 46 | -56.14 |
| 1MEQA | 23 | 44 | 69.95 |
| 1MKCA | 43 | 93 | -161.84 |
| 1MM0A | 36 | 59 | -306.42 |
| 1MNTB | 66 | 120 | 1.3 |
| 1MR0A | 34 | 62 | -334.69 |
| 1NAUA | 27 | 55 | -21.72 |
| 1NCSA | 47 | 86 | -40.64 |
| 1ND9A | 49 | 125 | -108.8 |
| 1NEIB | 60 | 112 | 65.68 |
| 1NG7B | 60 | 112 | 23.9 |
| 1NMJA | 28 | 59 | -2.37 |
| 1NWDB | 28 | 48 | -31.69 |
| 1NYBA | 22 | 40 | -24.26 |
| 1NZ9A | 58 | 89 | 89.91 |
| 1O9AB | 24 | 26 | 1.44 |
| 1ODPA | 20 | 37 | -84.43 |
| 1OEFA | 24 | 41 | 3.69 |
| 1OIGA | 24 | 40 | -163.66 |

| | | | |
|---|---|---|---|
| 1ONVB | 21 | 40 | -30.84 |
| 1OZSB | 20 | 27 | 18.8 |
| 1P1PA | 24 | 36 | -297.77 |
| 1P83A | 25 | 55 | -7.49 |
| 1P8BA | 37 | 57 | -77.86 |
| 1P9DS | 32 | 60 | 80.59 |
| 1PB5A | 35 | 77 | -271.29 |
| 1PD7B | 24 | 40 | -68.83 |
| 1PEHA | 33 | 48 | -83.76 |
| 1PEIA | 22 | 45 | -72.45 |
| 1PFSB | 78 | 123 | 108.49 |
| 1PLPA | 25 | 45 | 2.29 |
| 1PLSA | 113 | 229 | -134.76 |
| 1PSVA | 28 | 58 | -58.8 |
| 1PXQA | 33 | 57 | 64.81 |
| 1Q0WA | 24 | 52 | -43.38 |
| 1Q1VA | 70 | 150 | -32.23 |
| 1Q2IA | 32 | 61 | -65.75 |
| 1Q3JA | 36 | 58 | -232.79 |
| 1Q68A | 38 | 59 | -158.15 |
| 1Q69B | 29 | 47 | -41.39 |
| 1Q8LA | 84 | 173 | -1.72 |
| 1QFNB | 25 | 28 | 9.89 |
| 1QG9A | 21 | 50 | 13.5 |
| 1QJLA | 28 | 42 | -192.87 |
| 1QLDA | 50 | 119 | -178.91 |
| 1QYPA | 57 | 92 | -33.33 |
| 1R7GA | 31 | 56 | -58.15 |
| 1R8UA | 50 | 75 | 33.13 |
| 1RETA | 43 | 96 | 15.39 |
| 1RI9A | 77 | 153 | -62.11 |
| 1RIJA | 23 | 45 | 60.61 |
| 1RIMA | 33 | 65 | -126.4 |
| 1RJIA | 31 | 58 | -233.07 |
| 1RPCA | 21 | 29 | -118.2 |
| 1S4ZC | 30 | 33 | 39.22 |
| 1S5RA | 23 | 43 | 50.96 |
| 1S6WA | 21 | 33 | -434.48 |
| 1S8KA | 29 | 54 | -278.58 |
| 1SG5A | 86 | 165 | -56.82 |
| 1SKHA | 30 | 57 | 6.89 |
| 1SMZA | 27 | 53 | -9.53 |
| 1SO9A | 131 | 236 | 181.22 |
| 1SOLA | 20 | 30 | -12.75 |
| 1SP7A | 24 | 39 | -122.22 |
| 1SPFA | 35 | 79 | -62.25 |
| 1SS3A | 50 | 100 | -473.15 |

| | | | |
|---|---|---|---|
| 1SSEA | 35 | 55 | -61.48 |
| 1SSEB | 86 | 144 | -127.88 |
| 1SSLA | 48 | 94 | -452.58 |
| 1SUTA | 22 | 47 | 86.01 |
| 1SZLA | 61 | 105 | -147.6 |
| 1T0CA | 31 | 54 | 111.69 |
| 1T1HA | 78 | 154 | -16.56 |
| 1T3OA | 84 | 146 | -57.78 |
| 1T50A | 58 | 121 | -357.13 |
| 1T8JA | 22 | 27 | -39.17 |
| 1TERA | 21 | 46 | -220.82 |
| 1TI5A | 46 | 113 | -523.44 |
| 1TM6A | 22 | 48 | -93.93 |
| 1TPNA | 50 | 76 | -223.6 |
| 1TXPD | 28 | 55 | -127.6 |
| 1U34A | 119 | 175 | -157.23 |
| 1UEOA | 63 | 89 | -24.6 |
| 1UG2A | 95 | 166 | -59.67 |
| 1UGLA | 50 | 104 | -399.25 |
| 1UJLA | 42 | 64 | 73.16 |
| 1USTA | 92 | 186 | 108.04 |
| 1UT3A | 38 | 56 | -222.87 |
| 1UUCA | 55 | 131 | -280.04 |
| 1V1DA | 31 | 65 | 55.23 |
| 1V32A | 101 | 218 | -62.59 |
| 1V5AA | 28 | 50 | -270.97 |
| 1V7FA | 29 | 54 | -326.28 |
| 1VD4A | 62 | 112 | -71.53 |
| 1VIBA | 54 | 125 | -618.67 |
| 1VRYA | 61 | 142 | 66.95 |
| 1VTPA | 26 | 57 | 36.94 |
| 1W2QA | 127 | 286 | -717.64 |
| 1WA7B | 22 | 29 | 109.31 |
| 1WEWA | 78 | 112 | 50.06 |
| 1WEXA | 104 | 188 | 61.96 |
| 1WEYA | 104 | 186 | -69.97 |
| 1WFTA | 123 | 173 | 146.14 |
| 1WIDA | 117 | 191 | 16.26 |
| 1WIJA | 127 | 241 | 44.11 |
| 1WIVA | 73 | 144 | 44.32 |
| 1WJ0A | 58 | 90 | -133.13 |
| 1WJUA | 100 | 168 | -9.81 |
| 1WLPA | 25 | 34 | 191.52 |
| 1WM8A | 28 | 40 | -166.65 |
| 1WN4A | 28 | 57 | -16.29 |
| 1WN8A | 22 | 43 | -24.89 |
| 1WNNA | 23 | 51 | -26.51 |

| | | | |
|---|---|---|---|
| 1WO6A | 25 | 43 | -85.1 |
| 1WPIA | 133 | 318 | -74.3 |
| 1WQBA | 32 | 50 | -316.96 |
| 1WQEA | 23 | 57 | -236.22 |
| 1WSOA | 32 | 64 | -168.33 |
| 1WXNA | 42 | 74 | -217.18 |
| 1WZ4A | 23 | 45 | 40.26 |
| 1X32A | 47 | 92 | 94.14 |
| 1X3UA | 79 | 196 | 121.75 |
| 1X4TA | 92 | 183 | -45.14 |
| 1X58A | 62 | 113 | -7.33 |
| 1X5JA | 113 | 172 | 201.15 |
| 1X5VA | 33 | 67 | -289.8 |
| 1XBDA | 87 | 141 | 9.41 |
| 1XI7A | 47 | 71 | -199.09 |
| 1XKMC | 22 | 42 | 50.45 |
| 1XKMD | 25 | 50 | -65.37 |
| 1XNLA | 28 | 51 | 63.14 |
| 1XO3A | 101 | 213 | -36.61 |
| 1XOOA | 20 | 39 | 47.39 |
| 1XOPA | 20 | 35 | 34.27 |
| 1XR0A | 22 | 28 | 5.94 |
| 1Y0JB | 36 | 59 | -45.47 |
| 1Y4EA | 26 | 46 | 72.61 |
| 1Y9JA | 140 | 327 | -86.72 |
| 1YTRA | 26 | 52 | -17.41 |
| 1YUJA | 54 | 101 | -89.75 |
| 1YWWA | 65 | 146 | -160.7 |
| 1YYBA | 26 | 50 | -35.22 |
| 1YZ2A | 26 | 53 | -349.89 |
| 1Z2TA | 23 | 38 | -13.31 |
| 1Z6WA | 49 | 73 | -147.5 |
| 1Z9IA | 53 | 101 | -58.51 |
| 1ZAQA | 44 | 70 | -185.09 |
| 1ZECA | 25 | 60 | 65.47 |
| 1ZFDA | 32 | 56 | -125.09 |
| 1ZFOA | 30 | 49 | -91.01 |
| 1ZJQA | 34 | 56 | -232.85 |
| 1ZL8B | 54 | 137 | -91.22 |
| 1ZPXA | 21 | 31 | -276.84 |
| 1ZRWA | 25 | 52 | -75.13 |
| 1ZRXA | 42 | 97 | 36.3 |
| 1ZSGB | 22 | 25 | 103.58 |
| 1ZTOA | 36 | 57 | -48.84 |
| 1ZUFA | 42 | 72 | -315.03 |
| 1ZW8A | 64 | 113 | -119.47 |
| 2A2BA | 41 | 78 | -0.32 |

| | | | |
|---|---|---|---|
| 2A7UA | 22 | 42 | -31.15 |
| 2A93B | 32 | 70 | -113.67 |
| 2AB7A | 29 | 50 | -85.64 |
| 2AB9A | 31 | 47 | 11.85 |
| 2AGHC | 31 | 43 | 90.61 |
| 2AJOA | 28 | 47 | -11.87 |
| 2AMNA | 26 | 47 | 39.49 |
| 2AP8A | 19 | 33 | 73.4 |
| 2B19A | 36 | 84 | -25.73 |
| 2B7EA | 59 | 127 | -30.05 |
| 2B9KA | 47 | 77 | -1.53 |
| 2B9ZB | 74 | 179 | 27.17 |
| 2BBGA | 40 | 79 | -588.31 |
| 2BBPA | 22 | 28 | 59.01 |
| 2BDSA | 43 | 79 | -208.73 |
| 2BI6H | 41 | 74 | -207.99 |
| 2BL6A | 37 | 62 | -97.07 |
| 2BN5B | 21 | 22 | 99.9 |
| 2BN6A | 33 | 77 | -28.78 |
| 2BZTA | 66 | 139 | -127.4 |
| 2CH0A | 133 | 210 | -166.71 |
| 2CO8A | 82 | 132 | -212.52 |
| 2CQNA | 77 | 148 | -60.82 |
| 2CR8A | 53 | 66 | -93.58 |
| 2CT5A | 73 | 122 | -53.03 |
| 2CT7A | 86 | 136 | -199.22 |
| 2CTTA | 104 | 140 | -69.53 |
| 2D35A | 62 | 100 | 151.49 |
| 2D46A | 61 | 152 | -152.31 |
| 2DA2A | 70 | 136 | -26.67 |
| 2DARA | 90 | 142 | -86.57 |
| 2DBGA | 103 | 205 | -76.29 |
| 2DCEA | 111 | 206 | -12.93 |
| 2DCOA | 34 | 77 | -82.25 |
| 2DCVA | 42 | 65 | -243.45 |
| 2DI0A | 71 | 167 | -23.19 |
| 2DI8A | 111 | 173 | 234.78 |
| 2DSMA | 72 | 111 | -188.25 |
| 2DW3A | 77 | 137 | 0.21 |
| 2DWFA | 34 | 66 | -155.14 |
| 2DY8A | 69 | 132 | -55.52 |
| 2E1XA | 27 | 47 | -128.43 |
| 2E2SA | 35 | 61 | -329.99 |
| 2E3EA | 45 | 80 | -338.43 |
| 2E60A | 101 | 185 | 21.06 |
| 2E6IA | 64 | 85 | -14.77 |
| 2EEMA | 34 | 68 | -486.68 |

| | | | |
|---|---|---|---|
| 2EKKA | 47 | 109 | -145.27 |
| 2ELVA | 36 | 69 | -112.28 |
| 2ESYA | 31 | 57 | 82.62 |
| 2EWLA | 56 | 103 | -274.58 |
| 2EZDA | 21 | 27 | 91.61 |
| 2EZHA | 65 | 145 | -86.44 |
| 2FC6A | 50 | 69 | 33.57 |
| 2FCGF | 25 | 43 | -1.2 |
| 2FGXA | 85 | 159 | -112.3 |
| 2FHWA | 24 | 50 | -106.7 |
| 2FLYA | 20 | 39 | -28.88 |
| 2FMCA | 82 | 130 | -213.62 |
| 2FMRA | 65 | 111 | -33.32 |
| 2FQCA | 25 | 57 | -128.61 |
| 2G2KA | 156 | 293 | 51.34 |
| 2G9PA | 26 | 60 | 52.54 |
| 2GD3A | 24 | 37 | 51.34 |
| 2GDLA | 31 | 56 | 42.16 |
| 2GL1A | 47 | 89 | -451.38 |
| 2GLGA | 32 | 68 | -105.56 |
| 2GLOA | 59 | 128 | -197.18 |
| 2GP8A | 40 | 106 | 3.39 |
| 2GUTA | 77 | 168 | -137.4 |
| 2GVSA | 109 | 205 | -303.35 |
| 2GX1A | 29 | 64 | -350.98 |
| 2H8BB | 31 | 57 | 0.31 |
| 2H9RC | 22 | 40 | -16.76 |
| 2HACA | 33 | 74 | -28.65 |
| 2HFQA | 85 | 157 | -51.83 |
| 2HGOA | 26 | 43 | -221.41 |
| 2HJJA | 66 | 129 | 14.02 |
| 2HJQA | 111 | 212 | -99.22 |
| 2HLGA | 39 | 72 | -354.95 |
| 2HN8A | 38 | 99 | -19.97 |
| 2HP8A | 68 | 178 | -463.25 |
| 2HTGA | 27 | 51 | -21.57 |
| 2I2JA | 21 | 43 | -0.52 |
| 2I9NA | 33 | 60 | 42.48 |
| 2IGRA | 33 | 79 | 7.02 |
| 2IKDA | 56 | 89 | -256.85 |
| 2IKEA | 54 | 94 | -198.71 |
| 2J5HA | 39 | 51 | -225.34 |
| 2J8PA | 49 | 110 | -64.58 |
| 2JMXB | 25 | 34 | 17.63 |
| 2JNIA | 21 | 35 | -69.07 |
| 2JNRB | 23 | 38 | 74.83 |
| 2JO4D | 20 | 38 | -15.76 |

| | | | |
|---|---|---|---|
| 2JODB | 33 | 56 | -21.47 |
| 2JOFA | 20 | 34 | 49.39 |
| 2JP6A | 37 | 62 | -267.86 |
| 2JPKA | 35 | 68 | 23.96 |
| 2JPXA | 23 | 51 | -31.79 |
| 2JQTA | 57 | 112 | -42.15 |
| 2JR3A | 42 | 70 | -197.4 |
| 2JRAA | 67 | 95 | -24.1 |
| 2JRRA | 67 | 119 | -225.48 |
| 2JRWA | 23 | 32 | 52.65 |
| 2JSHA | 23 | 41 | 30.33 |
| 2JTBA | 33 | 49 | -245.07 |
| 2JTGA | 87 | 175 | -21 |
| 2JTTC | 31 | 57 | 7.41 |
| 2JU5A | 144 | 269 | -111.96 |
| 2JUCA | 55 | 142 | -118.29 |
| 2JWHA | 48 | 75 | -235.25 |
| 2JWKA | 74 | 142 | 45.33 |
| 2JX1A | 31 | 51 | -25 |
| 2JX8A | 47 | 84 | -16.96 |
| 2JX9A | 106 | 191 | -54.03 |
| 2JXCB | 40 | 63 | 66.36 |
| 2JXFA | 30 | 61 | -19.96 |
| 2JY0A | 27 | 44 | 21.86 |
| 2JYAA | 80 | 125 | 91.78 |
| 2JYPA | 36 | 86 | -213.9 |
| 2JYVA | 32 | 44 | -122.31 |
| 2JZ7A | 81 | 122 | 75.66 |
| 2JZ8A | 87 | 140 | -142.63 |
| 2JZIB | 24 | 64 | 2.91 |
| 2K1OA | 54 | 83 | -75.8 |
| 2K1VA | 20 | 36 | -118.64 |
| 2K29A | 50 | 94 | 64.24 |
| 2K2DA | 47 | 67 | -34.94 |
| 2K2IB | 20 | 35 | -71.83 |
| 2K2SB | 55 | 97 | -75.61 |
| 2K2UB | 35 | 49 | 64.13 |
| 2K38A | 35 | 55 | -2.98 |
| 2K3CA | 31 | 54 | 46.08 |
| 2K3JA | 65 | 120 | -109.35 |
| 2K3XA | 115 | 207 | 11.32 |
| 2K44A | 28 | 48 | -4.13 |
| 2K4FA | 57 | 97 | 219.58 |
| 2K4UA | 37 | 53 | -202.3 |
| 2K58B | 35 | 61 | -24.44 |
| 2K5DA | 110 | 177 | 57.03 |
| 2K6LA | 51 | 93 | 67.79 |

| | | | |
|---|---|---|---|
| 2K6UA | 27 | 53 | -129.65 |
| 2K72A | 37 | 81 | -230.21 |
| 2K7RA | 106 | 218 | -186.84 |
| 2K8EA | 116 | 215 | -17.3 |
| 2K8FB | 39 | 64 | 99.82 |
| 2K8JX | 29 | 50 | 68.3 |
| 2K98A | 24 | 48 | 32.61 |
| 2K9BA | 33 | 67 | -33.88 |
| 2K9HA | 57 | 100 | -254.14 |
| 2K9UB | 24 | 27 | 69.56 |
| 2KA4B | 57 | 90 | 192.52 |
| 2KA6B | 45 | 67 | 90.39 |
| 2KAMA | 25 | 54 | -36.85 |
| 2KBCB | 24 | 47 | -13.99 |
| 2KBIA | 97 | 160 | 71.33 |
| 2KBVA | 26 | 54 | 39.75 |
| 2KC1A | 91 | 173 | 165.62 |
| 2KD3A | 97 | 186 | -133 |
| 2KDTA | 43 | 71 | -34.48 |
| 2KDUB | 36 | 77 | -23.39 |
| 2KELA | 46 | 95 | -150.52 |
| 2KETA | 26 | 57 | 28.4 |
| 2KEYA | 112 | 216 | -173.92 |
| 2KFEA | 24 | 51 | -13.76 |
| 2KGNA | 24 | 48 | -3.49 |
| 2KGRA | 111 | 221 | 147.56 |
| 2KHFA | 25 | 57 | 27.93 |
| 2KHSB | 35 | 71 | -127.57 |
| 2KHWA | 32 | 60 | 14.94 |
| 2KI9A | 33 | 74 | -58.51 |
| 2KIOA | 33 | 69 | -100.42 |
| 2KIXA | 33 | 74 | -11.98 |
| 2KJEB | 42 | 61 | 112.97 |
| 2KJIA | 50 | 114 | -408.06 |
| 2KJNA | 26 | 58 | -136.34 |
| 2KJXA | 65 | 121 | -154.59 |
| 2KJZA | 122 | 216 | 184.22 |
| 2KKTA | 52 | 108 | -59.87 |
| 2KL7A | 71 | 109 | 49.21 |
| 2KM9A | 25 | 50 | -267.6 |
| 2KNNA | 29 | 45 | -186.94 |
| 2KNSA | 33 | 64 | 48.55 |
| 2KNUA | 29 | 57 | 48.3 |
| 2KOZA | 33 | 65 | -289.91 |
| 2KPAA | 26 | 50 | 19.72 |
| 2KQCA | 47 | 62 | 70.83 |
| 2KQSB | 20 | 19 | -0.4 |

| | | | |
|---|---|---|---|
| 2KR7A | 151 | 212 | 39.19 |
| 2KSEA | 77 | 149 | 3.02 |
| 2KSFA | 107 | 241 | 126.26 |
| 2KSWA | 66 | 125 | -181.27 |
| 2KV5A | 33 | 68 | 7.47 |
| 2KVFA | 28 | 55 | -59.03 |
| 2KVGA | 27 | 49 | -24.16 |
| 2KVHA | 27 | 54 | -91.7 |
| 2KVLA | 61 | 139 | 20.14 |
| 2KVRA | 128 | 244 | -176.16 |
| 2KVXA | 28 | 46 | -262.52 |
| 2KW0A | 81 | 172 | -120.26 |
| 2KWOB | 19 | 19 | 44.32 |
| 2KWTA | 33 | 52 | -13.29 |
| 2KWVA | 36 | 63 | 115.74 |
| 2KWZA | 40 | 71 | -2.65 |
| 2KXQB | 20 | 22 | 95.77 |
| 2KYAA | 34 | 63 | 36.67 |
| 2KYGC | 38 | 68 | -38.99 |
| 2KYJA | 36 | 61 | -105.56 |
| 2KZ0A | 76 | 134 | 3.47 |
| 2KZBA | 114 | 216 | 140.79 |
| 2KZQA | 36 | 71 | 57.99 |
| 2KZYA | 62 | 88 | -1.83 |
| 2L0EA | 31 | 64 | 55.73 |
| 2L0NA | 30 | 48 | -20.43 |
| 2L0YB | 21 | 37 | -41.86 |
| 2L0ZA | 41 | 66 | -47.77 |
| 2L1OA | 37 | 61 | -4.39 |
| 2L1WB | 25 | 55 | -11.88 |
| 2L2IB | 27 | 36 | 14.53 |
| 2L2LB | 36 | 73 | -105.21 |
| 2L2RA | 37 | 70 | -256.35 |
| 2L3IA | 30 | 62 | -121.75 |
| 2L58A | 32 | 65 | -52.11 |
| 2L5BA | 31 | 61 | 16.4 |
| 2L5JA | 20 | 27 | 75.97 |
| 2L5RA | 23 | 45 | 47.37 |
| 2L63A | 33 | 65 | -39.22 |
| 2L6SA | 20 | 20 | 59.03 |
| 2L6WA | 39 | 78 | 8.7 |
| 2L79A | 39 | 77 | 26.63 |
| 2L7SA | 52 | 79 | -62.26 |
| 2L91A | 43 | 97 | 112.99 |
| 2L9BA | 91 | 194 | 74.57 |
| 2L9UA | 40 | 78 | -61.83 |
| 2L9ZA | 39 | 56 | -32.05 |

| | | | |
|---|---|---|---|
| 2LA0A | 29 | 63 | -65.09 |
| 2LA2A | 37 | 74 | 52.15 |
| 2LAIA | 101 | 154 | 100.55 |
| 2LAUA | 81 | 160 | -132.48 |
| 2LBGA | 27 | 55 | 113.45 |
| 2LCMA | 28 | 64 | 10.86 |
| 2LCVA | 47 | 93 | 46.6 |
| 2LCYA | 54 | 105 | 40.97 |
| 2LDJA | 19 | 27 | 41.44 |
| 2LE3A | 42 | 67 | 4.46 |
| 2LE7A | 20 | 31 | -23.53 |
| 2LE8B | 29 | 55 | -71.5 |
| 2LEHB | 26 | 42 | -6.47 |
| 2LFWB | 62 | 89 | 44.35 |
| 2LG5A | 36 | 71 | -308.2 |
| 2LH9A | 78 | 150 | 79.94 |
| 2LHUA | 35 | 71 | 99.41 |
| 2LI3A | 30 | 51 | -312.06 |
| 2LI5B | 34 | 51 | 10.45 |
| 2LI8A | 63 | 104 | -140.21 |
| 2LICA | 34 | 39 | -9.44 |
| 2LIFA | 27 | 53 | 36.93 |
| 2LIXA | 27 | 47 | -234.27 |
| 2LIYA | 45 | 76 | -224.58 |
| 2LJTA | 37 | 69 | 74.88 |
| 2LJYA | 70 | 125 | -239.53 |
| 2LJZA | 75 | 176 | -250.83 |
| 2LK0A | 30 | 48 | -173.38 |
| 2LK9A | 24 | 61 | 17.27 |
| 2LKEA | 24 | 45 | 73.94 |
| 2LKWA | 20 | 22 | 77.26 |
| 2LLLA | 121 | 209 | 13.17 |
| 2LLMA | 43 | 93 | 66.09 |
| 2LLRA | 22 | 31 | -141.33 |
| 2LLWA | 71 | 160 | 144.99 |
| 2LLZA | 100 | 214 | 25.56 |
| 2LMAA | 22 | 43 | 130.73 |
| 2LMFA | 23 | 52 | -34.02 |
| 2LMJA | 66 | 113 | -2.03 |
| 2LN8A | 75 | 170 | -495.69 |
| 2LNHC | 47 | 63 | 118.84 |
| 2LNYA | 20 | 29 | -0.25 |
| 2LO2A | 38 | 75 | -133.36 |
| 2LO4A | 28 | 52 | -30.98 |
| 2LOCA | 23 | 49 | -511.6 |
| 2LONA | 99 | 199 | 93.11 |
| 2LOXB | 20 | 22 | 30.73 |

| | | | |
|---|---|---|---|
| 2LP0B | 20 | 28 | -22.48 |
| 2LPBB | 34 | 54 | -22.73 |
| 2LQ0A | 25 | 46 | -22.3 |
| 2LQ1A | 25 | 46 | 87.64 |
| 2LQ6A | 79 | 129 | -81.97 |
| 2LQCB | 24 | 43 | -31.33 |
| 2LQLA | 83 | 148 | -403.11 |
| 2LQXA | 41 | 92 | -380.92 |
| 2LQYA | 20 | 37 | -8.04 |
| 2LR1B | 21 | 33 | -72.6 |
| 2LR5A | 38 | 79 | -254.46 |
| 2LR7A | 29 | 54 | -96.6 |
| 2LRDA | 61 | 144 | -587.23 |
| 2LS1A | 20 | 27 | -126.37 |
| 2LS2A | 25 | 51 | 51.24 |
| 2LS3A | 29 | 73 | -52.34 |
| 2LS4A | 24 | 39 | 27.34 |
| 2LS9A | 25 | 47 | 46.38 |
| 2LSQA | 25 | 41 | -193.28 |
| 2LSWA | 35 | 67 | 44.05 |
| 2LTUA | 62 | 108 | -55.68 |
| 2LU2A | 81 | 185 | 76.66 |
| 2LUFA | 20 | 41 | 81.82 |
| 2LV2A | 85 | 128 | -266.67 |
| 2LVHA | 45 | 86 | -127.37 |
| 2LVUA | 26 | 46 | -86.35 |
| 2LWBA | 20 | 40 | -188.28 |
| 2LWRA | 68 | 104 | -175.94 |
| 2LX0A | 32 | 72 | -26.72 |
| 2LXRA | 76 | 152 | -6.45 |
| 2LXTC | 33 | 59 | -39.82 |
| 2LYDA | 134 | 245 | -64.56 |
| 2LYDB | 38 | 49 | 81.19 |
| 2LYWA | 24 | 28 | -0.72 |
| 2LYXA | 87 | 133 | -52.55 |
| 2LZPA | 31 | 51 | 69.7 |
| 2LZQA | 26 | 50 | -82.26 |
| 2LZRA | 48 | 95 | 41.98 |
| 2LZXA | 36 | 72 | -215.15 |
| 2M0EA | 29 | 48 | -113.78 |
| 2M0HA | 22 | 38 | -32.76 |
| 2M0KB | 28 | 55 | -12.83 |
| 2M0WA | 23 | 52 | 30.22 |
| 2M1JA | 30 | 52 | -27.39 |
| 2M59A | 37 | 86 | -51.53 |
| 2M5LA | 20 | 23 | 90.71 |
| 2M5ZA | 43 | 88 | -5.36 |

| | | | |
|---|---|---|---|
| 2M6AA | 28 | 58 | -399.19 |
| 2M6JA | 25 | 38 | -143.17 |
| 2M6NA | 46 | 74 | -230.49 |
| 2M6OA | 48 | 79 | 7.46 |
| 2M6QA | 91 | 162 | -81.76 |
| 2M71A | 98 | 193 | -87.45 |
| 2M73A | 25 | 47 | -63.64 |
| 2M7EA | 26 | 46 | -26 |
| 2M8FA | 24 | 46 | 51.96 |
| 2M8SB | 24 | 34 | -22.84 |
| 2M9HA | 106 | 166 | 19 |
| 2MAEA | 25 | 51 | -42.13 |
| 2MAKD | 23 | 49 | -82.72 |
| 2MC7A | 30 | 53 | -83.2 |
| 2MCEA | 21 | 44 | 3.93 |
| 2MCQA | 77 | 129 | -34.37 |
| 2MCRA | 36 | 81 | -272.51 |
| 2MCUA | 22 | 46 | 13.91 |
| 2MD0A | 51 | 108 | -322.11 |
| 2MDLA | 24 | 29 | -69.06 |
| 2MDUA | 29 | 54 | -38.29 |
| 2MF3A | 47 | 62 | -217.53 |
| 2MFKA | 69 | 118 | -85.15 |
| 2MFLA | 93 | 158 | 76.94 |
| 2MFPA | 31 | 53 | -140.36 |
| 2MFQB | 22 | 38 | 120.22 |
| 2MG1A | 27 | 48 | 22.37 |
| 2MG2A | 28 | 44 | -21 |
| 2MGPA | 99 | 210 | -197.15 |
| 2MGUM | 36 | 59 | -43.09 |
| 2MH0A | 39 | 63 | 64.53 |
| 2MH3A | 70 | 150 | -160.92 |
| 2MH4A | 92 | 149 | 94.94 |
| 2MHEA | 74 | 124 | 50.34 |
| 2MI5A | 57 | 88 | -292.35 |
| 2MI9A | 30 | 48 | -156.48 |
| 2MICA | 41 | 80 | -17.67 |
| 2MIJA | 37 | 74 | -195.16 |
| 2MIXA | 21 | 36 | -344.94 |
| 2MIZA | 156 | 276 | -9.08 |
| 2MJ2A | 36 | 68 | -98 |
| 2MJCA | 24 | 31 | -44.55 |
| 2MJKA | 41 | 83 | -213.1 |
| 2MK9A | 34 | 69 | -16.31 |
| 2MKBA | 25 | 43 | 4.64 |
| 2MKCC | 33 | 46 | 45.69 |
| 2MKDA | 60 | 92 | -84.66 |

| | | | |
|---|---|---|---|
| 2MLPA | 26 | 54 | 20.18 |
| 2MLUA | 30 | 52 | 61.48 |
| 2MM0A | 64 | 115 | -65.09 |
| 2MMLA | 70 | 109 | -7.09 |
| 2MMUA | 50 | 94 | -2.68 |
| 2MN8A | 20 | 38 | 19.62 |
| 2MNIA | 92 | 185 | 219.35 |
| 2MNJA | 23 | 24 | 93.8 |
| 2MNSA | 20 | 36 | 53.01 |
| 2MNUB | 26 | 34 | -23.42 |
| 2MP2C | 25 | 24 | 9.24 |
| 2MPCA | 90 | 190 | 2.23 |
| 2MPJA | 25 | 38 | -32.05 |
| 2MQKA | 65 | 132 | -62.77 |
| 2MQMA | 118 | 191 | -126.62 |
| 2MQUA | 35 | 61 | -279.06 |
| 2MRBA | 31 | 41 | -157.64 |
| 2MSOA | 30 | 55 | -461.65 |
| 2MSRA | 21 | 21 | 22.4 |
| 2MSUA | 20 | 34 | -8.88 |
| 2MTGA | 116 | 223 | 129.79 |
| 2MTPB | 21 | 38 | 81.17 |
| 2MTWA | 20 | 21 | -27.31 |
| 2MTXA | 20 | 22 | 24.7 |
| 2MTYA | 20 | 25 | 12.73 |
| 2MU6A | 20 | 25 | -7.22 |
| 2MU7A | 20 | 25 | -19.85 |
| 2MU9A | 20 | 31 | -43.52 |
| 2MUAA | 21 | 27 | 1.75 |
| 2MUDA | 20 | 34 | -15.01 |
| 2MUEA | 20 | 33 | 24.11 |
| 2MUFA | 20 | 26 | -29.62 |
| 2MUGA | 20 | 25 | -16.39 |
| 2MUJA | 21 | 29 | -48.48 |
| 2MV3A | 88 | 204 | -155.05 |
| 2MV7B | 25 | 32 | 61.62 |
| 2MVAA | 27 | 43 | -119.01 |
| 2MVIA | 43 | 59 | -89.79 |
| 2MVJA | 26 | 52 | 25.03 |
| 2MVTA | 47 | 94 | -392.91 |
| 2MW3A | 21 | 33 | 78.29 |
| 2MW7A | 37 | 67 | -220.79 |
| 2MWNB | 93 | 191 | -24.4 |
| 2MX2A | 81 | 170 | 54.07 |
| 2MX4A | 43 | 63 | 83.11 |
| 2MY3B | 23 | 30 | 39.98 |
| 2MYQA | 43 | 87 | -10.11 |

| | | | |
|---|---|---|---|
| 2MYYA | 80 | 135 | 56.86 |
| 2MZ0A | 55 | 106 | -249.91 |
| 2MZFA | 64 | 96 | -379.66 |
| 2MZPI | 27 | 43 | -24.89 |
| 2MZRA | 95 | 150 | 42.76 |
| 2N01A | 23 | 29 | 69.11 |
| 2N1AB | 53 | 84 | -146.64 |
| 2N1EA | 19 | 22 | 20.38 |
| 2N1PA | 30 | 55 | 7.87 |
| 2N1UA | 70 | 125 | -189.68 |
| 2N21A | 20 | 33 | 27.41 |
| 2N24A | 30 | 54 | 26.3 |
| 2N25A | 29 | 50 | -46 |
| 2N2HA | 27 | 47 | -29.7 |
| 2N2QA | 54 | 93 | -405.65 |
| 2N2SA | 32 | 62 | -489.85 |
| 2N30A | 23 | 34 | -15.51 |
| 2N31A | 22 | 29 | 67.4 |
| 2N34A | 70 | 91 | 26.79 |
| 2N39A | 108 | 215 | -231.58 |
| 2N4FA | 84 | 145 | 50.13 |
| 2N4NA | 20 | 25 | -55.2 |
| 2N4QA | 24 | 31 | 105.28 |
| 2N55B | 40 | 50 | 38.87 |
| 2N58A | 30 | 58 | -50.15 |
| 2N5KA | 29 | 44 | -67.57 |
| 2N5MA | 52 | 122 | -139.73 |
| 2N5QA | 27 | 54 | -414.94 |
| 2N5UA | 79 | 131 | -65.39 |
| 2N68A | 23 | 35 | 51.94 |
| 2N6BA | 34 | 68 | -204.75 |
| 2N6MA | 21 | 41 | 12.65 |
| 2N6NA | 32 | 52 | -341.7 |
| 2N6UA | 20 | 33 | 0.69 |
| 2N77B | 33 | 66 | 70.41 |
| 2N7FA | 32 | 49 | -267.7 |
| 2N7IA | 37 | 79 | -16.54 |
| 2N7QA | 46 | 103 | 27.01 |
| 2N86A | 59 | 91 | -441.94 |
| 2N8DA | 20 | 35 | 6.56 |
| 2N8EA | 33 | 60 | -277.26 |
| 2N8HA | 34 | 49 | -265.93 |
| 2N8JB | 22 | 30 | 18.79 |
| 2N8NA | 72 | 129 | 44.23 |
| 2N8OA | 51 | 111 | 13.28 |
| 2N90A | 39 | 73 | -15.83 |
| 2N92A | 31 | 49 | 72.58 |

| | | | |
|---|---|---|---|
| 2N9CA | 20 | 31 | 46.34 |
| 2N9OA | 42 | 69 | -113.91 |
| 2N9YA | 42 | 82 | -14.27 |
| 2NA6A | 28 | 58 | -33.26 |
| 2NA7A | 28 | 51 | -5.61 |
| 2NA8A | 44 | 92 | -49.19 |
| 2NABA | 41 | 75 | -81.01 |
| 2NAEA | 43 | 58 | 109.28 |
| 2NATA | 28 | 52 | -34.89 |
| 2NAVA | 28 | 51 | -193.57 |
| 2NB0A | 55 | 100 | -307 |
| 2NB2A | 38 | 87 | -376.08 |
| 2NC2A | 56 | 101 | -425.93 |
| 2NCXA | 24 | 59 | -23.09 |
| 2NDDA | 25 | 47 | -254 |
| 2NDIA | 46 | 63 | -251.44 |
| 2NR1A | 23 | 56 | 42.48 |
| 2NSVA | 52 | 107 | -442.79 |
| 2NVJA | 25 | 43 | -89.05 |
| 2NX6A | 27 | 51 | -276.9 |
| 2NX7A | 28 | 61 | -340.41 |
| 2NZZA | 37 | 74 | -120.16 |
| 2OFQA | 95 | 150 | 48.02 |
| 2OQ9A | 24 | 40 | -114.07 |
| 2ORUA | 20 | 33 | 2.05 |
| 2PCOA | 26 | 47 | -40.43 |
| 2PFUA | 99 | 166 | 154.24 |
| 2PTAA | 35 | 69 | -192.09 |
| 2RLWA | 34 | 79 | 148.54 |
| 2RMRA | 71 | 133 | 99.41 |
| 2RMSB | 61 | 99 | 75.57 |
| 2RMYA | 34 | 70 | -23.74 |
| 2RODB | 27 | 50 | 57.74 |
| 2ROOA | 43 | 84 | -626.87 |
| 2RPJA | 50 | 85 | -193.58 |
| 2RPSA | 32 | 46 | 45.42 |
| 2RPWX | 25 | 47 | -34.03 |
| 2RQ2A | 30 | 56 | 99.41 |
| 2RQHA | 22 | 27 | 91.38 |
| 2RQWA | 105 | 198 | 97.47 |
| 2RQWB | 24 | 25 | 106.86 |
| 2RR3B | 44 | 48 | 24.82 |
| 2RREA | 74 | 145 | -19.32 |
| 2RSYB | 37 | 58 | 96.36 |
| 2RT4A | 25 | 39 | 63.49 |
| 2RTSA | 73 | 128 | 104.9 |
| 2RU1A | 70 | 112 | -333.29 |

| | | | |
|---|---|---|---|
| 2RVBA | 48 | 75 | -86.58 |
| 2VXDA | 54 | 115 | -8.93 |
| 2VY4A | 37 | 68 | -27.7 |
| 2W0TA | 43 | 79 | 10.65 |
| 2W84B | 20 | 38 | -10.03 |
| 2XC7A | 104 | 230 | -110.53 |
| 2XK0A | 69 | 119 | 42.12 |
| 2Y4QA | 79 | 170 | -77.64 |
| 2YKAB | 23 | 41 | -16.57 |
| 2YOMA | 23 | 47 | -31.67 |
| 2YONA | 29 | 69 | -49.82 |
| 2YRAA | 74 | 111 | -93.79 |
| 2YRTA | 75 | 141 | -70.04 |
| 2YT5A | 66 | 111 | -122.46 |
| 2YUCA | 76 | 119 | 0.9 |
| 2YUSA | 79 | 148 | -90.57 |
| 3MRAA | 25 | 61 | -1.64 |
| 3ZJ1A | 83 | 128 | -87.91 |
| 3ZNFA | 30 | 57 | -63.34 |
| 4A54B | 52 | 69 | 62.57 |
| 4ASVA | 79 | 168 | 27.3 |
| 4B19A | 30 | 62 | -1.46 |
| 4B2VA | 32 | 54 | -211.46 |
| 4BXLC | 22 | 32 | 47.74 |
| 4UZMA | 119 | 240 | 94.47 |
| 5B88A | 85 | 144 | 115.74 |
| 5FZVA | 32 | 64 | -321.18 |
| 5GWGB | 31 | 52 | -153.58 |
| 5GWMB | 50 | 119 | -106.01 |
| 5HUZB | 44 | 88 | -196.14 |
| 5IJ4A | 49 | 86 | -258.99 |
| 5IONA | 29 | 58 | -103.01 |
| 5IX5A | 68 | 135 | -236.35 |
| 5J6ZA | 30 | 39 | 119.59 |
| 5J8TA | 47 | 73 | -72.21 |
| 5JI4A | 37 | 72 | -278.78 |
| 5JN6A | 70 | 122 | -20.35 |
| 5JR0A | 34 | 62 | -63.82 |
| 5JTME | 25 | 28 | 50.56 |
| 5JTPE | 22 | 26 | -2.19 |
| 5JYHA | 44 | 71 | -216.85 |
| 5KHBA | 20 | 37 | 16.76 |
| 5KJGA | 22 | 29 | 21.32 |
| 5KVNA | 26 | 40 | -189.08 |
| 5KWXA | 24 | 40 | -111.9 |
| 5KWZA | 26 | 61 | -231.69 |
| 5KX1A | 22 | 55 | -1.41 |

| | | | |
|---|---|---|---|
| 5L1CA | 36 | 66 | -340.45 |
| 5L85B | 34 | 70 | -52.74 |
| 5LBJA | 35 | 70 | 25.6 |
| 5LCBA | 59 | 94 | 103.76 |
| 5LGMA | 69 | 146 | -4.55 |
| 5LV6A | 44 | 81 | 69.24 |
| 5LWCA | 49 | 105 | -35.77 |
| 5LXLA | 81 | 137 | 187.6 |
| 5M9ZA | 56 | 90 | -16.45 |
| 5MGQA | 36 | 76 | -110.77 |
| 5MPGA | 97 | 187 | -1.57 |
| 5NAMA | 48 | 104 | 44.72 |
| 5NPGA | 83 | 150 | 11.82 |
| 5T7QA | 21 | 42 | 1.93 |
| 5TP6B | 29 | 43 | 21.28 |
| 5U9RA | 23 | 48 | 35.24 |
| 5V2GA | 20 | 48 | -48.31 |
| 5VFWA | 25 | 35 | -33.09 |
| 5W54A | 25 | 30 | -4.78 |
| 5WCVA | 29 | 52 | -264.5 |
| 5X39A | 36 | 65 | -105.99 |
| 5X3LA | 20 | 40 | 50.67 |
| 5XIVA | 20 | 35 | -506.22 |
| 5XJKA | 65 | 125 | 133.67 |
| 5Y0IA | 21 | 33 | -149.35 |
| 8TFVA | 21 | 34 | -44.34 |
